# The *Firre* locus produces a trans-acting RNA molecule that functions in hematopoiesis

**DOI:** 10.1101/648279

**Authors:** Jordan P. Lewandowski, James C. Lee, Taeyoung Hwang, Hongjae Sunwoo, Jill M. Goldstein, Abigail F. Groff, Nydia Chang, William Mallard, Adam Williams, Jorge Henao-Meija, Richard A. Flavell, Jeannie T. Lee, Chiara Gerhardinger, Amy J. Wagers, John L. Rinn

## Abstract

RNA has been classically known to play central roles in biology, including maintaining telomeres^1^, protein synthesis^2^, and in sex chromosome compensation in certain species^3,4^. At the center of these important biological systems are noncoding RNAs. While thousands of long noncoding RNAs (lncRNAs) have been identified in mammalian genomes^5–8^, attributing RNA-based roles to lncRNA loci requires an assessment of whether the observed effect could be due to DNA regulatory elements, the act of transcription, or the lncRNA transcript. Here, we use the syntenically conserved lncRNA locus, <u>F</u>unctional intergenic repeating <u>R</u>NA <u>e</u>lement (*Firre*), that is located on the X chromosome as a model to discriminate between DNA- and RNA-mediated effects *in vivo*. To this end, we generated genetically defined loss-of-function, gain-of-function, and rescue mouse models for *Firre* and provide genetic evidence that the *Firre* locus produces a *trans*-acting RNA. We report that: (i) *Firre* mutant mice have cell-specific defects during hematopoiesis and changes in gene expression that can be rescued by induction of *Firre* RNA from a transgene in the *Firre* knockout background, (ii) mice overexpressing *Firre* from a transgene exhibit increased levels of pro-inflammatory cytokines and impaired survival upon exposure to lipopolysaccharide, and (iii) deletion of the *Firre* locus did not result in changes in local gene expression on the X chromosome in 9 different biological contexts, suggesting that *Firre* does not function by *cis*-acting RNA or DNA elements. Together, our results provide genetic evidence that the *Firre* locus produces a *trans*-acting lncRNA that has physiological roles in hematopoiesis and immune function.

## INTRODUCTION

Transcription occurs at thousands of sites throughout the mammalian genome. Many of these sites are devoid of protein-coding genes and instead contain long noncoding RNAs (lncRNAs). While lncRNA loci have been implicated in a variety of biological functions, comparatively few lncRNA loci have been genetically defined to have RNA-based roles. Indeed, deletions of entire lncRNA loci have uncovered a number of *in vivo* phenotypes^9–13^; however, this approach alone is confounded because in addition to the lncRNA transcript, lncRNA loci can also exert function through DNA regulatory elements^14–16^, the promoter region^17^, as well as by the act of transcription^18,19^. Thus, attributing RNA-based role(s) to lncRNA loci requires testing whether other regulatory modes potentially present at the locus have molecular activity that could contribute to phenotypic effects^11,20,21^.

In this study, we use the *Firre* locus as a model to discriminate between DNA- and RNA-mediated effects *in vivo*. We selected this locus for our study because it is syntenically conserved in a number of mammals including human^22–25^, and because studies have reported diverse biological and molecular roles. Early characterization of the *FIRRE* locus in human cell lines identified it as a region that interacts with the X-linked macrosatellite region, *DXZ4*, in a CTCF-dependent manner^26–29^. Further analyses of the *Firre* locus demonstrated that it produces a lncRNA that escapes X-inactivation^23,30–32^, although it is predominately expressed from the active X chromosome^29^. Studies using cell culture models suggest that the *Firre* locus has biological roles in multiple processes, including adipogenesis^33^, nuclear architecture^23,27,29^, and regulation of gene expression programs^23,34^. Additionally, there is some evidence for roles in human development and disease^35–38^. Collectively, these studies demonstrate the diverse cellular and biological functions for the *Firre* locus. However, the biological roles of *Firre* as well as disentangling DNA- and RNA-mediated function(s) for the *Firre* locus have not been explored *in vivo*.

Using multiple genetic approaches, we describe an *in vivo* role for the *Firre* locus during hematopoiesis. We report that *Firre* mutant mice have cell-specific defects in hematopoietic populations. Deletion of *Firre* is accompanied by significant changes in gene expression in a hematopoietic progenitor cell type, which can be rescued by induction of *Firre* RNA from a transgene within the *Firre* knockout background. Mice overexpressing *Firre* have increased levels of pro-inflammatory cytokines and impaired survival upon exposure to lipopolysaccharide (LPS). Finally, the *Firre* locus does not contain *cis*-acting RNA or DNA elements (including the promoter) that regulate neighboring gene expression on the X chromosome (9 biological contexts examined), suggesting that *Firre* does not function in *cis*. Collectively, our study provides evidence for a *trans*-acting RNA-based role for the *Firre* locus that has physiological importance for hematopoiesis and immune function.

## RESULTS

### The *Firre* locus produces an abundant lncRNA

We first sought to investigate the gene expression properties for *Firre* RNA *in vivo*. To determine potential spatial and temporal aspects of *Firre* RNA expression during development, we performed *in situ* hybridization in wildtype (WT) mouse embryos (E8.0 – E12.5). Notably, we detected *Firre* RNA in many embryonic tissues, including the forebrain, midbrain, pre-somitic mesoderm, lung, forelimb, hindlimb, liver, and heart (Fig. 1A). Since noncoding RNAs have been described to be generally expressed at lower levels compared to protein-coding genes^39–42^, we determined the relative abundance of *Firre* RNA *in vivo*. We performed RNA-seq on eight different WT embryonic tissues and plotted the expression of noncoding and coding transcripts. Consistent with previous reports^39–42^, we observed that noncoding transcripts were generally less abundant than protein-coding transcripts (Fig. 1B). Despite most lncRNAs being expressed at low levels, we found that *Firre*, like *Malat1*^43–45^, is an abundant transcript (Fig. 1B). Next, since *Firre* is located on the X chromosome and escapes X-inactivation^23,30–32^, we investigated whether *Firre* has different expression levels in male and female WT tissues. While levels of *Firre* RNA varied across embryonic tissue types, within individual tissues, male and female samples exhibited similar expression levels of *Firre*, despite escaping X inactivation (Fig. 1C).

**Figure 1.**
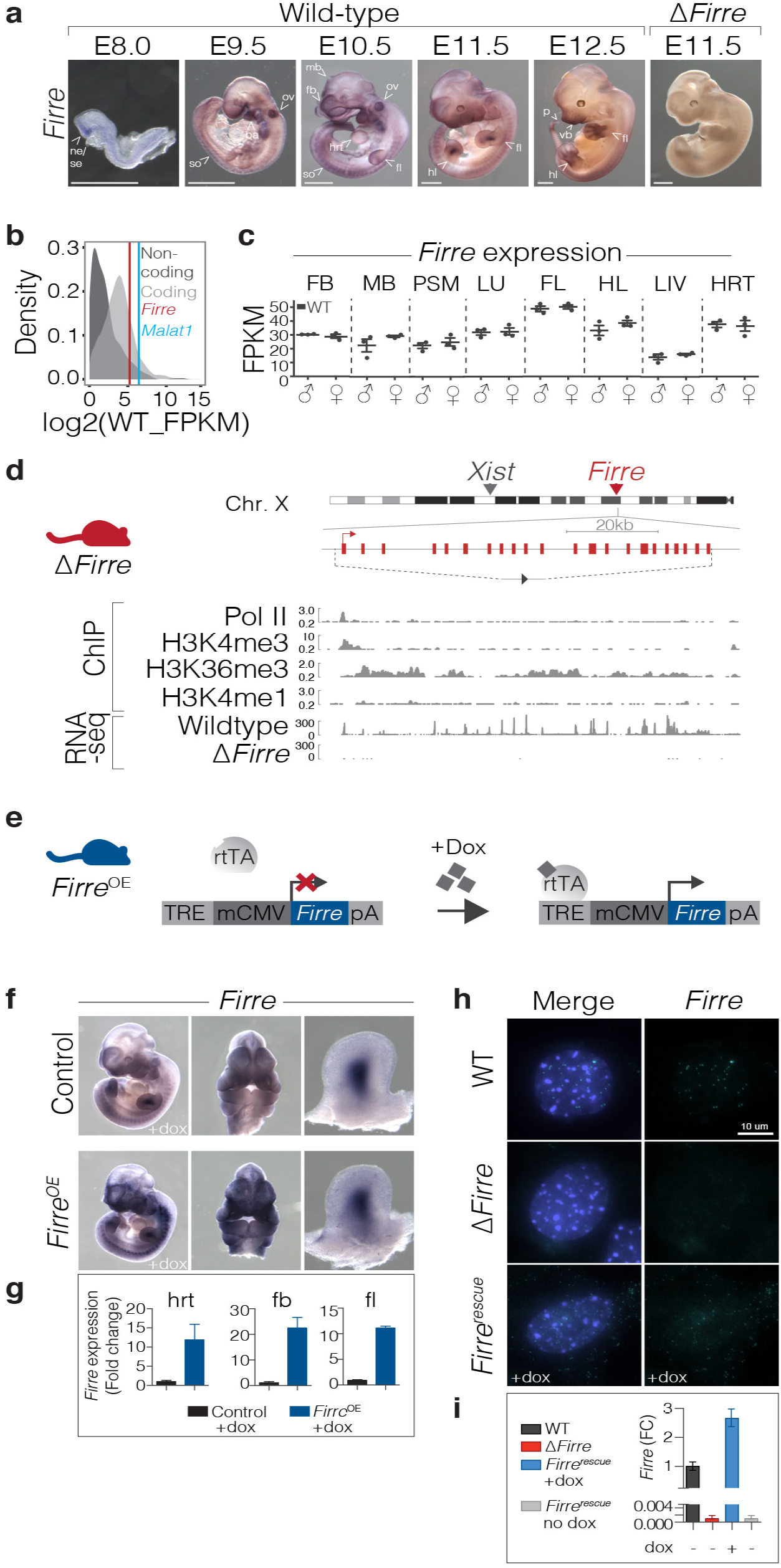
Mouse models to interrogate the *in vivo* function of *Firre*. **(A)** Whole-mount *in situ* hybridization for *Firre* RNA in WT mouse embryos at E8.0 (*n*=4), E9.5 (*n*=4), E10.5 (*n*=5), E11.5 (*n*=6), and E12.5 (*n*=4) and Δ*Firre* E11.5 embryos (*n*=3). Scale bar is equal to 1 mm. **(B)** Abundance for protein coding transcripts (light gray) and noncoding transcripts (dark gray) in WT E11.5 heart tissue (representative tissue shown from 7 additional tissues). Vertical lines indicate *Firre* (red) and *Malat1* (blue). **(C)** Expression of *Firre* in E11.5 WT male (*n*=3) and female (*n*=3) tissues shown as fragments per kilobase of transcript per million mapped reads (FPKM) from RNA-seq. Data shown as mean ± standard error of the mean (SEM). Tissue abbreviations: forebrain (FB), midbrain (MB), pre-somitic mesoderm (PSM), lung (LU), forelimb (FL), hindlimb (HL), liver (LIV), and heart (HRT). **(D)** *Firre* knockout mouse (red). Schematic of mouse X chromosome ideogram showing the *Firre* locus relative to *Xist*. UCSC genome browser diagram of the *Firre* locus (shown in opposite orientation). Dashed lines indicate the genomic region that is deleted in Δ*Firre* mice; single loxP scar upon deletion (gray triangle). Histone modifications and transcription factor binding sites in mouse embryonic stem cells (mESC-Bruce4, ENCODE/LICR, mm9). RNA-seq tracks for the *Firre* locus in WT and Δ*Firre* E11.5 forelimbs. **(E)** Schematic of doxycycline(dox)-inducible *Firre* overexpression mouse (*Firre*^*OE*^, blue). Tet-responsive element (TRE), minimal CMV promoter (mCMV), reverse-tetracycline transcriptional activator (rtTA), beta-globin polyA terminator (pA). **(F)** *in situ* hybridization for *Firre* at E11.5 in control (WT or tg(*Firre*) +dox) (n=4) and *Firre*^*OE*^ +dox (*n*=3) embryos. **(G)** qRT-PCR for *Firre* expression shown as fold-change (FC) in dox-treated E11.5 control and *Firre*^*OE*^ hrt, fb, and fl. Expression normalized to beta-actin in the control sample and data plotted as mean ± confidence interval (CI) at 98%. **(H)** RNA-FISH for *Firre* in male WT, Δ*Firre*, and *Firre*^rescue^ MEFs. DAPI (blue) marks the nucleus and *Firre* RNA is shown in green. **(I)** qRT-PCR for *Firre* expression shown as FC in male WT, Δ*Firre, Firre*^rescue^ +dox, and *Firre*^rescue^ no dox MEFs. Expression normalized to beta-actin in the WT sample and data plotted as mean ± CI at 98%.

### *Firre* knockout and overexpression mice are viable and fertile

To investigate the *in vivo* role of *Firre* and assess DNA- and RNA-mediated effects, we generated both *Firre* loss-of-function and *Firre* overexpression mice. To delete the *Firre* locus *in vivo*, we generated a mouse line containing a floxed allele (*Firre*^floxed^) from a previously targeted mouse embryonic stem cell line^23^ and matted to a CMV-Cre deleter mouse^46^. This produced a genomic deletion (81.8 kb) that removed the entire *Firre* gene body and promoter (henceforth called Δ*Firre*) (Fig. 1D). We confirmed the deletion of the *Firre* locus by genotyping (Extended Data Fig. 1) and examined *Firre* RNA expression. As expected, we did not detect *Firre* RNA in Δ*Firre* embryos by whole-mount *in situ* hybridization or by RNA-seq (Fig. 1A, D).

Since *Firre* is found on the X chromosome, we first sought to determine if deletion of the locus had an effect on the expected sex ratio of the progeny. Matings between Δ*Firre* mice produced viable progeny with a normal frequency of male and female pups that did not exhibit overt morphological, skeletal, or weight defects (Extended Data Table 1 and Extended Data Fig. 2). Moreover, deletion of *Firre* did not impact expression levels of *Xist* RNA in embryonic tissues or perturb *Xist* RNA localization during random X chromosome inactivation (XCI) in mouse embryonic fibroblasts (MEFs) (Extended Data Fig. 3A-C).

Because the Δ*Firre* allele removes the entire gene body, this model does not allow us to distinguish between DNA- and RNA-mediated effects. Therefore, in order to be able to investigate the role of *Firre* RNA, we generated a doxycycline (dox)-inducible *Firre* overexpression mouse. This mouse model was engineered to contain a *Firre* cDNA downstream of a tet-responsive element (henceforth called tg(*Firre*)) and was mated to mice that constitutively express the reverse tetracycline transcriptional activator (*rtTA*) gene (combined alleles henceforth called *Firre*^OE^) (Fig. 1E). This approach enabled systemic induction of *Firre* RNA in a temporally controllable manner by the administration of dox. Moreover, by combining the *Firre*^OE^ and Δ*Firre* alleles (henceforth called *Firre*^rescue^) we could test whether *Firre* RNA expression alone is sufficient to rescue phenotypes arising in the Δ*Firre* mice, thereby distinguishing DNA- and RNA-based effects.

To confirm expression of transgenic *Firre* RNA, tg(*Firre*) females were mated with *rtTA* males and placed on a dox diet the day a copulatory plug was detected, and embryos were collected at E11.5 for analyses. Compared to sibling control embryos, we detected increased *Firre* RNA in *Firre*^OE^ embryos by whole-mount *in situ* hybridization (Fig. 1F) and by quantitative reverse transcription-PCR (qRT-PCR) (heart, 16 fold; forebrain, 26.6 fold; and forelimb, 11.5 fold) (Fig. 1G). Moreover, mice overexpressing *Firre* are viable and detected at expected male and female frequencies (Extended Data Table 1).

*Firre* RNA has been reported to be largely enriched in the nucleus of mouse embryonic stem cells (mESCs)^23,47^, neuronal precursor cells^39^, and HEK293 cells^17^, but also has been reported in the cytoplasm of a human colon cell line^34^. Thus, we investigated the subcellular localization of *Firre* in the genetic models using RNA fluorescent *in situ* hybridization (RNA FISH). In contrast to Δ*Firre* MEFs, we detected pronounced localization of *Firre* RNA in the nucleus of WT MEFs (Fig. 1H). In dox-treated *Firre*^rescue^ MEFs, which only produce *Firre* RNA from the transgene, we detected *Firre* RNA in both the nucleus and cytoplasm (Fig. 1H), which corresponded to approximately a 2.7-fold increase in *Firre* RNA relative to WT (Fig. 1I). Notably, the *Firre*^rescue^ transgenic model showed both nuclear and cytoplasmic localization of *Firre*, suggesting a threshold level control for nuclear localized *Firre*.

### RNA-seq of Δ*Firre* embryonic tissues identifies tissue-specific gene dysregulation

Given the broad expression profile of *Firre* RNA (Fig. 1A), we took an initial unbiased approach to explore the potential biological roles for the *Firre* locus, and performed poly(A)+ RNA-seq on eight E11.5 tissues from WT and Δ*Firre* embryos (forebrain, midbrain, heart, lung, liver, forelimb, hindlimb, and pre-somitic mesoderm). As expected, *Firre* expression was not detected in any of the Δ*Firre* tissues (Extended Data Tables 2-9). Deletion of *Firre* was accompanied by significant changes in gene expression in all tissues examined (>1FPKM, FDR<0.05) (Fig. 2A,B and Extended Data Tables 2-9).

**Figure 2.**
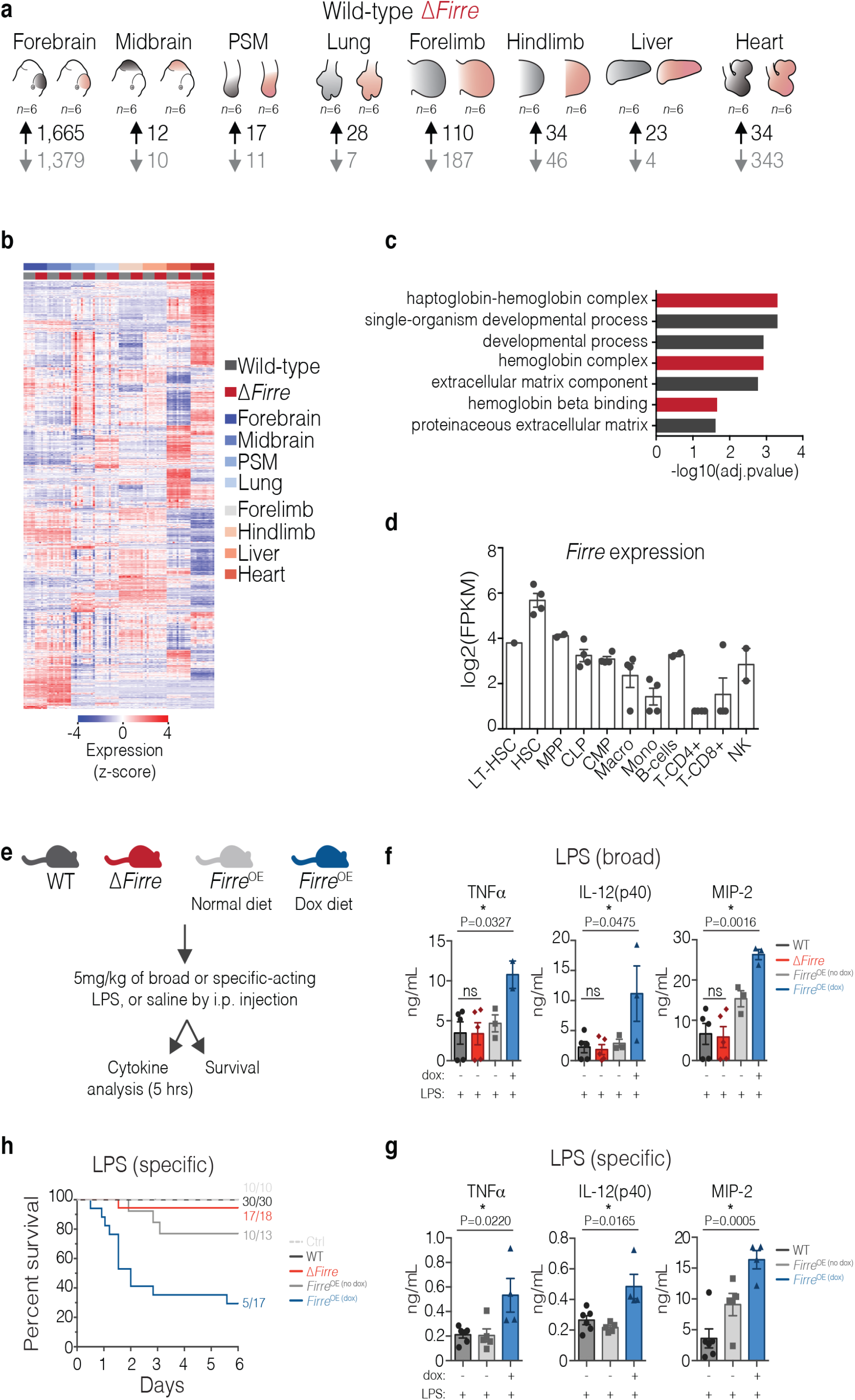
Modulation of *Firre* impacts genes with roles in the blood. **(A)** Schematized E11.5 tissues used for RNA-seq. WT (*n*=6) shown in black and Δ*Firre* (*n*=6) shown in red. Number of differentially expressed genes shown below each tissue. **(B)** Heatmap of replicate embryonic tissues. **(C)** GO analysis for genes found dysregulated in four or more tissues. **(D)** *Firre* expression across multiple mouse blood cell lineages (RNA-seq data from bloodspot.eu, GSE60101). **(E)** Experimental approach for cytokine and survival experiments. **(F)** Cytokine measurements in serum at 5 hours post intraperitoneal (i.p.) injection of 5 mg/kg LPS (broad-acting) in WT (*n*=5), Δ*Firre* (*n*=5), *Firre*^OE^ control diet (*n*=3), and *Firre*^OE^ dox diet (*n*=2 to 3). Data are shown as mean ± SEM and significance determined by an unpaired two-tail t-test. **(G)** Cytokine measurements in serum at 5 hours post i.p. injection of 5mg/kg LPS (specific-acting) in WT (*n*=6), *Firre*^OE^ control diet (*n*=5), *Firre*^OE^ dox diet (*n*=4). Data are shown as mean ± SEM and significance determined by an unpaired two-tail t-test. **(H)** 6-day survival plot of mice injected with 5 mg/kg LPS (specific-acting) or saline over two independent experiments in WT (*n*=30), Δ*Firre* (*n*=18), *Firre*^OE^ control diet (*n*=13), and *Firre*^OE^ dox diet (*n*=17). Saline control group (*n*=10) consisting of WT, Δ*Firre*, and *Firre*^OE^ mice. Significance determined by Mantel-Cox test.

Across these eight tissues, we identified a total of 3,910 significantly differentially expressed genes, of which 271 genes were differentially expressed in two or more tissues (Extended Data Tables 2-9). Interestingly, gene ontology (GO) analysis of the commonly dysregulated genes showed that deletion of the *Firre* locus affected genes involved in hemoglobin regulation and general blood developmental processes (Fig. 2C). We therefore analyzed publicly available mouse RNA-seq datasets and found that *Firre* is expressed across many blood cell types and note that expression is found highest in hematopoietic stem cells (HSCs)^48^ and then decreases in conjunction with hematopoietic differentiation^49^ (Fig. 2D). Based on this information we narrowed our investigation to evaluate potential roles for *Firre* in the blood system, and leveraged the genetic mouse models to test DNA- and RNA-mediated effects.

### LPS exposure to mice overexpressing *Firre* RNA impacts the innate immune response

*Firre* is expressed in many innate immune cell types (Fig. 2D) and has been shown to regulate the levels of inflammatory genes in human intestinal epithelial and mouse macrophage cell lines^34^. Thus, we hypothesized that dysregulation of *Firre* might alter the inflammatory response *in vivo*. To test this, we employed a commonly used endotoxic shock model by administering lipopolysaccharide (LPS) intraperitoneally to cohorts of WT, Δ*Firre, Firre*^OE^ no dox, and dox-fed *Firre*^OE^ mice in order to stimulate signaling pathways that regulate inflammatory mediators^50^ (Fig. 3E).

**Figure 3.**
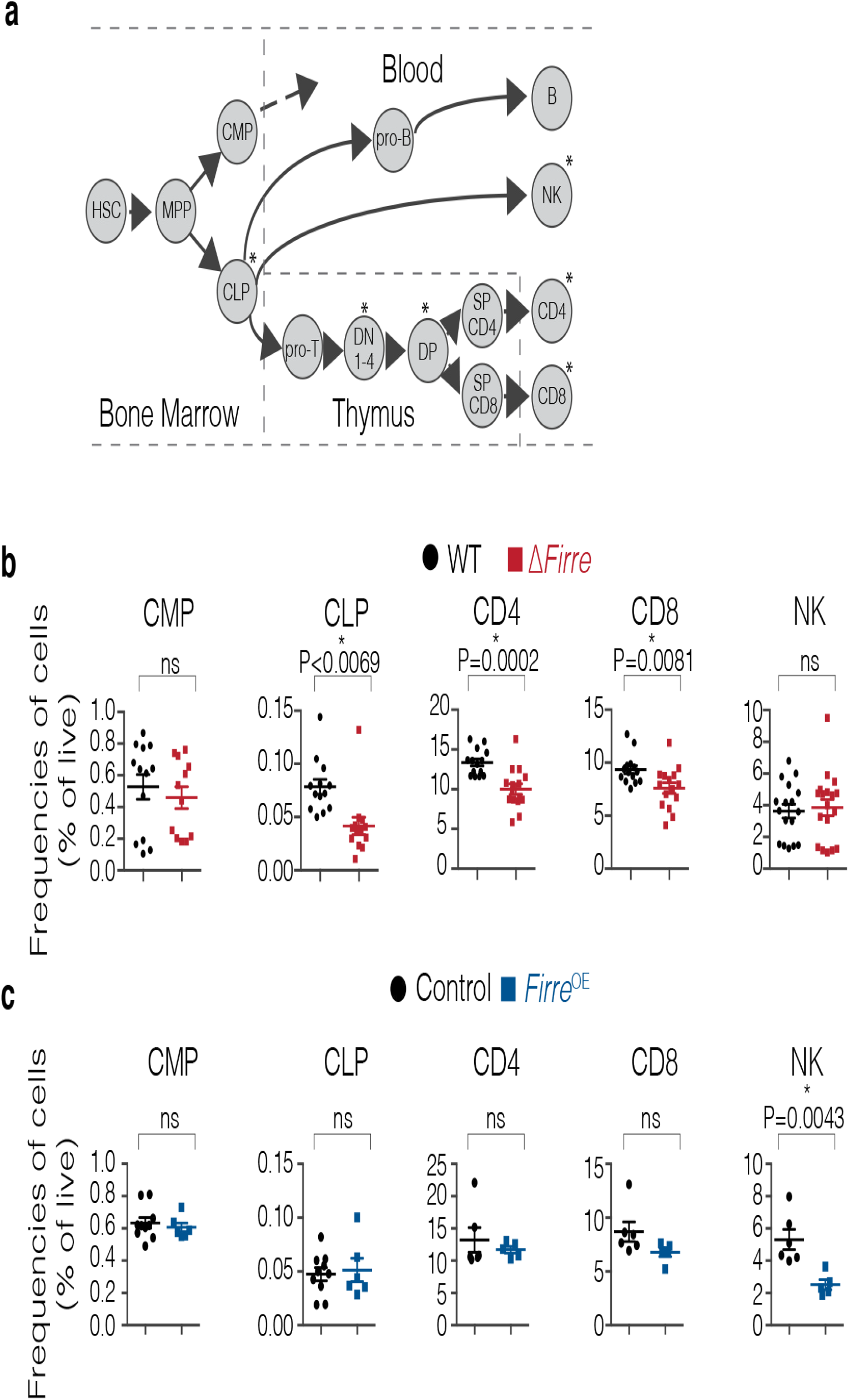
Δ*Firre* and *Firre*^OE^ mice have cell-specific defects during hematopoiesis. **(A)** Schematic of hematopoiesis. **(B)** Frequencies of CD4, CD8, and NK cells from the peripheral blood from WT (black circle) and Δ*Firre* (red square) mice. Three representative experiments combined (seven independent experiments). Frequencies of common myeloid progenitors (CMP) and common lymphoid progenitors (CLP) in the bone marrow shown from WT and Δ*Firre* mice. Two representative experiments combined (three independent experiments). **(C)** Frequencies of CMPs and CLPs from the bone marrow from control (tg(*Firre*) or WT or rtTA with dox) (black circle) and dox-treated *Firre*^OE^ (blue square) mice. One representative experiment shown (two independent experiments). Frequencies of CD4, CD8, and NK cells from the peripheral blood from control (WT or tg(*Firre*) or rtTA with dox) (black circle) and dox-treated *Firre*^OE^ (blue square) mice. One representative experiment shown (three independent experiments). All cell frequencies determined by flow cytometry analysis. All data are plotted as percent (%) of live cells showing the mean ± SEM and statistical significance determined by a two-tailed Mann-Whitney U test.

We administered two different LPS preparations, one which broadly stimulates the pattern recognition receptors toll-like receptors (TLR) 2, 4 and nitric oxide synthase, and an ultrapure LPS preparation that specifically stimulates TLR4^51–53^. At 5 hours post LPS injection we measured serum cytokine levels. Notably, we observed that *Firre*^OE^ dox-fed mice administered broad-acting LPS had significantly higher levels of inflammatory cytokines, including TNFα, IL12-p40, and MIP-2 compared to WT (Fig. 2F). In contrast, we did not observe a significant difference for these cytokines in LPS-treated Δ*Firre* mice (Fig. 2F). Consistent with the increased cytokine response using broad-acting LPS, dox-fed *Firre*^OE^ mice administered TLR4 specific-acting LPS also had significantly higher levels of TNFα, IL12-p40, and MIP-2 compared to WT (Fig. 2G), albeit at lower serum concentrations compared to the broad-acting LPS (Fig. 2F,G). In addition, we confirmed that overexpressing *Firre* RNA alone (without LPS) does not result in increased serum levels of TNFα, IL12-p40, and MIP-2 (Extended Data Fig. 4).

Because increased levels of TNFα is a hallmark of endotoxic shock^54–56^, we next tested whether the levels of *Firre* RNA had an impact on survival following LPS treatment. We administered 5mg/kg of TLR4 specific-acting LPS to WT (*n*=30), Δ*Firre* (*n*=18), *Firre*^OE^ no dox (*n*=13), and *Firre*^OE^ dox-fed (*n*=17) mice, as well as a saline control group and monitored for 6 days. At this dose, across two independent cohorts, dox-treated *Firre*^OE^ mice showed a significantly higher susceptibility to LPS compared to WT mice (P<0.0001, Mantel-Cox) and uninduced *Firre*^OE^ animals (P=0.0063, Mantel-Cox) (Fig. 2H). Whereas Δ*Firre* mice did not show a significant difference in the level of susceptibility to LPS (P=0.1967, Mantel-Cox test) (Fig. 2H). Collectively, these results indicate that increased levels of *Firre* RNA can modulate the inflammatory response *in vivo* independent of genomic context, suggesting an RNA-based role for *Firre*.

### Modulating the *Firre* locus and RNA results in cell-specific defects during hematopoiesis

Having observed an effect of *Firre* in regulating gene expression and accentuating the inflammatory response, we further investigated the role of *Firre* in hematopoiesis (Fig. 3A). We first examined cell populations in the peripheral blood in Δ*Firre* mice and observed a modest but significant reduction in the frequencies of CD4 and CD8 T cells (P=0.0002 and P=0.0081, respectively), whereas the frequencies of B and NK cells were unaffected compared to WT (Fig. 3B and Extended Data Fig. 5A). To investigate the cause of this reduction we examined the thymus (to assess for a defect in T cell development) and the bone marrow (to assess for a defect in hematopoietic progenitor cells). There was no block in thymic development in Δ*Firre* mice, as normal frequencies of cells were observed at each developmental stage (Extended Data Fig. 5B, upper panels). However, we noticed that the absolute number of cells was generally lower in Δ*Firre* mice at every developmental stage, suggestive of a pre-thymic defect in progenitor development (Extended Data Fig. 5B, lower panels). Consistent with this, in the bone marrow compartment, we observed a significant reduction in both the frequency and number of the common lymphoid progenitors (CLPs) (lineage(lin)^-^Sca-1^lo^-c-Kit^lo^IL7Rα+), a hematopoietic progenitor cell type, in Δ*Firre* mice (P=0.0069 and P<0.0001, respectively) (Fig. 3B and Extended Data Fig. 5C).

To assess whether the observed defect in hematopoiesis could be due to a progenitor-intrinsic effect of *Firre* deficiency, we performed competitive chimera assays. Briefly, we isolated an HSC-enriched population (lin-Sca-1^+^c-Kit^+^CD34^+/-^CD135^-^) from WT (CD45.2) and Δ*Firre* (CD45.2), mixed the cells separately at an equal ratio with congenic WT (CD45.1) HSCs, and transplanted this mixture into lethally irradiated CD45.1 recipient mice (Extended Data Fig. 6A,B). We assessed the long-term reconstitution ability of WT and Δ*Firre* HSCs to repopulate blood cell lineages *in vivo*. We observed that Δ*Firre*/CD45.2-donors had reduced frequencies of CD4 and CD8 T-cells (*n*=10, P=0.0028 and P=0.0051), B-cell (*n*=10, P=0.0114), and NK cell (*n*=10, 0.0068) populations in the peripheral blood of recipient mice compared to WT/CD45.2-donors, suggesting that Δ*Firre*-donors were markedly outcompeted at repopulating the blood (Extended Data Fig. 6C). These data are consistent with a progenitor-intrinsic role for *Firre* in hematopoiesis.

In contrast to the Δ*Firre* model, mice overexpressing *Firre* RNA in the WT background (*Firre*^OE^), had normal frequencies of CD4, CD8 and B cells, but had a significant reduction in the frequency of NK cells (P=0.0043) in the peripheral blood compared to control mice (Fig. 3C, Extended Data Fig. 5D). A decrease in the frequency of NK cells in dox-fed *Firre*^rescue^ mice, where only *Firre* RNA from the transgene is expressed, was also observed (Extended Data Fig. 7). In the bone marrow of dox-treated *Firre*^OE^ mice, we did not observe significant changes in the frequencies of HSC, multipotent progenitor (MPP), common myeloid progenitor (CMP), or CLPs compared to control samples (Fig. 3C and Extended Data Fig. 5E). Taken together, these results identify cell type-specific defects during hematopoiesis, whereby alterations of *Firre* impact the ratios and numbers of particular blood cells produced during hematopoiesis.

### *Firre* lncRNA has a *trans*-acting role *in vivo*

Next, we wanted to further investigate the DNA- and RNA-mediated effects of the *Firre* locus using a cell type that was dysregulated in the Δ*Firre* immunophenotyping analysis. We selected to use the CLP as a model because this was the earliest hematopoietic defect identified and because *Firre* is highly expressed in this progenitor cell type. Therefore, further investigation could provide insight into the physiological effects of modulating *Firre* in a progenitor cell population. Because the Δ*Firre* mouse contains a deletion that removes all potential DNA-regulatory elements, the lncRNA, and the promoter (thus removing the act of transcription), this mouse model does not allow us to distinguish between DNA- and RNA-mediated effects.

To directly test whether the hematopoietic CLP defect in the Δ*Firre* mice is mediated by an RNA-based mechanism, we reasoned that overexpressing *Firre* RNA in the Δ*Firre* background would enable us to identify RNA-mediated effects. To this end, we generated multiple cohorts of compound mice (*Firre*^rescue^) that contained the *Firre*^OE^ alleles in the Δ*Firre* background and induced transgenic *Firre* expression by placing mice on a dox-diet. From multiple cohorts of WT, Δ*Firre*, and dox-fed *Firre*^rescue^ mice, we assessed CLP frequency by flow cytometry in total bone marrow and lineage depleted bone marrow (to enrich for hematopoietic non-lineage committed cells) (Fig. 4A).

**Figure 4.**
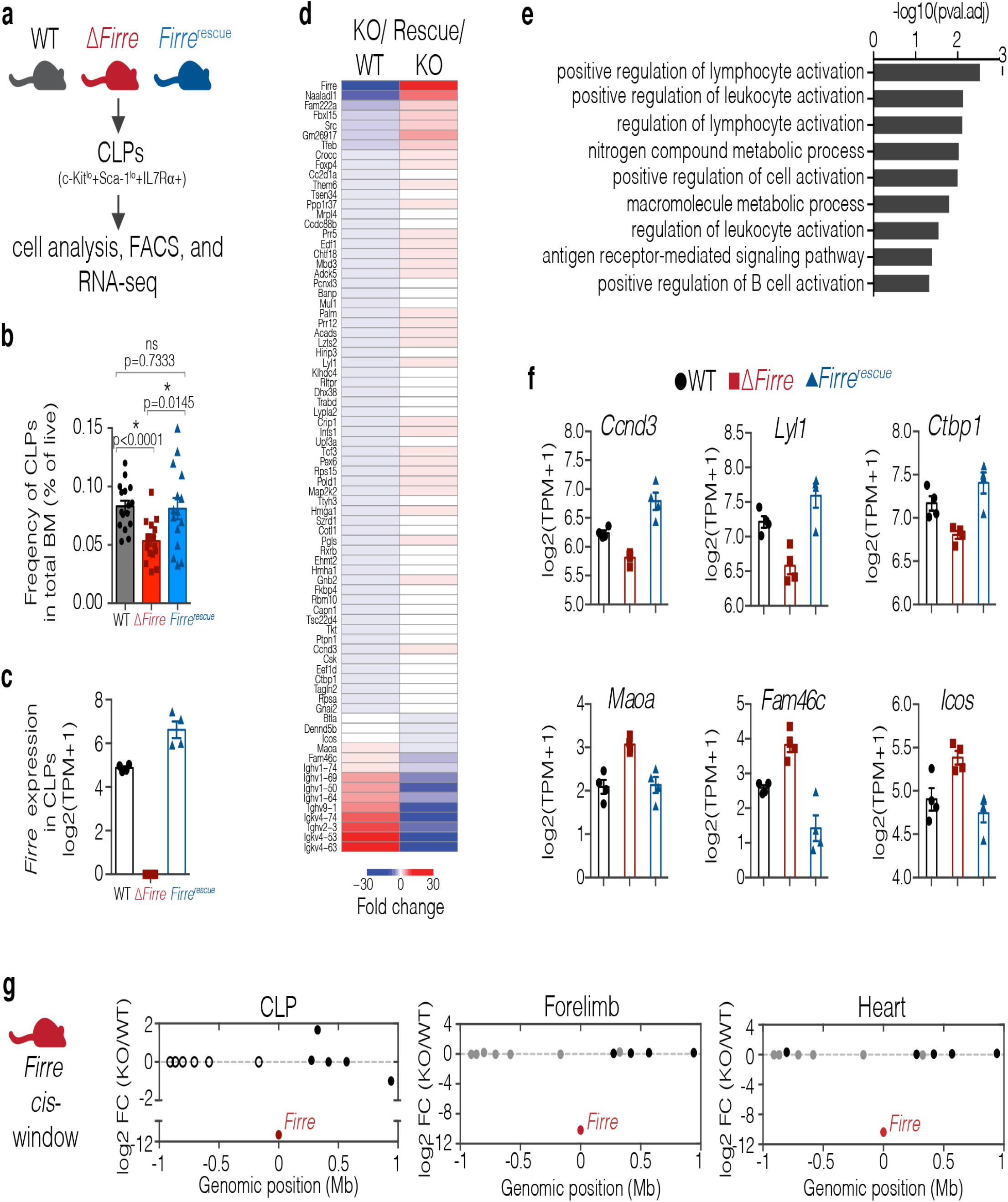
Ectopic expression of *Firre* rescues physiological and molecular defects in CLPs *in vivo*. **(A)** Schematic of experimental approach. **(B)** Bar graph indicating the frequency of CLPs shown as percent of live in total bone marrow from 3-7 month old WT (*n*=16, mean age=26 weeks), Δ*Firre* (*n*=17, mean age=23 weeks), and *Firre*^rescue^ dox diet (*n*=15, mean age=23 weeks) mice over three independent experiments. Data are shown as mean ± SEM and statistical significance determined by a two-tailed Mann-Whitney U test. **(C)** *Firre* RNA expression in CLPs from WT (*n*=4), Δ*Firre* (*n*=4), and dox-treated *Firre*^rescue^ (*n*=4) determined by RNA-seq. Data plotted as transcripts per million (TPM +1) showing the mean ± SEM. **(D)** Heatmap showing significantly differentially expressed genes in CLPs in Δ*Firre* / WT comparison and dox-treated *Firre*^rescue^ / Δ*Firre* and comparison. **(E)** GO analysis for significantly dysregulated genes in Δ*Firre* CLPs. **(F)** Examples of genes that show significant reciprocal regulation in WT, Δ*Firre*, and dox-treated *Firre*^rescue^ CLPs. **(G)** *Firre* locus region (2 Mb) showing gene expression differences in log2 FC between Δ*Firre* and WT CLPs, mouse embryonic forelimb, and heart. *Firre* is shown in red, significantly dysregulated genes are shown in red, genes that are not significantly changed are shown in black, and genes that were not detected shown in white.

Consistent with our previous data (Fig. 2B), we observed a significant decrease in the frequency of CLPs in total bone marrow from Δ*Firre* mice (*n*=17, mean CLP frequency = 0.0532) compared to WT mice (*n*=16, mean CLP frequency = 0.0829) (P<0.0001) (Fig. 4B). Further, in separate experimental cohorts, the frequency of CLPs in lineage-depleted bone marrow from Δ*Firre* mice was significantly decreased (*n*=9, mean CLP frequency = 0.3011, P=0.0071) compared to WT (*n*=9, mean CLP frequency = 0.2167) (Extended Data Fig. 8). Notably, the frequency of CLPs was significantly increased in dox-fed *Firre*^rescue^ mice compared to Δ*Firre* mice and restored to approximately that of WT in both total bone marrow (*n*=15, mean CLP frequency = 0.0810, P=0.0145) (Fig. 4B) and lineage depleted bone marrow (*n*=11, mean CLP frequency = 0.2809, P=0.0234, respectively) (Extended Data Fig. 8). Thus, induction of transgenic *Firre* RNA alone is sufficient to rescue the reduction in frequency of CLPs observed in Δ*Firre* bone marrow. These data suggest that *Firre* RNA, rather than the DNA, exerts a biological function in the CLPs during hematopoiesis.

### Expression of transgenic *Firre* restores gene expression programs *in vivo*

To gain further insight into the molecular roles of *Firre* in the CLPs, we took a gene expression approach because alterations of the *Firre* locus and RNA have previously been shown to impact gene expression^23,29,57^. Moreover, we reasoned that we could test if changes in gene expression in the loss-of-function model could be rescued by expressing only *Firre* RNA. To this end, we isolated CLPs by fluorescence activated cell sorting (FACS) from the bone marrow of age- and sex-matched WT, Δ*Firre*, and dox-fed *Firre*^rescue^ mice and performed poly(A)+ RNA-seq. As expected, *Firre* RNA was not detected in the Δ*Firre* samples, and *Firre* RNA levels were restored to levels above WT in the *Firre*^rescue^ samples (Fig. 4C). Differential gene expression analysis between WT and Δ*Firre* CLPs identified 89 significantly differentially expressed genes (FDR<0.1) (Fig. 4D and Extended Data Table 10). GO analysis of the differentially expressed genes showed that deletion of *Firre* in CLPs affected genes involved in lymphocyte activation, cell adhesion, and B cell activation (Fig. 4E).

Next, we determined if induction of *Firre* RNA in the *Firre*^rescue^ model could rescue expression of the 89 significantly dysregulated genes found in Δ*Firre* CLPs. We compared the CLP RNA-seq from Δ*Firre* and *Firre*^rescue^ mice and identified 4,656 genes with significant changes in gene expression (FDR<0.1) (Extended Data Table 11). Notably, 78 of the 89 genes that were significantly differentially expressed in Δ*Firre* CLPs, were found to be significantly and reciprocally regulated in *Firre*^rescue^ CLPs (P=2.2e-16, Fisher exact test) (Fig. 4D). For example, *Ccnd3, Lyl1*, and *Ctbp1* are significantly downregulated in Δ*Firre* CLPs, but are found significantly upregulated in *Firre*^rescue^ CLPs to (Fig. 4F). Further, genes such as *Maoa, Fam46c*, and *Icos* were found significantly upregulated in Δ*Firre* CLPs, but their expression was significantly reduced in *Firre*^rescue^ CLPs (Fig. 4F). We also noted that several immunoglobin heavy and light chain variable region genes were reciprocally regulated in our analyses (Fig. 4D). Taken together, these data suggest that ectopic expression of *Firre* is sufficient to restore a gene expression program in an RNA-based manner *in vivo*.

### *Firre* does not function in cis

Many lncRNA loci exert function to control the expression of neighboring genes, a biological function called *cis* regulation^58^. This occurs through a variety of mechanisms including, *cis*-acting DNA regulatory elements, the promoter region, the act of transcription, and the lncRNA (a biological function called *cis* regulation)^18,59–62^. The Δ*Firre* mouse model enables to test for a potential *cis* regulatory roles for *Firre* on the X chromosome because the knockout removes the entire *Firre* locus (Fig. 1D). To investigate local (*cis*) effects on gene expression, we generated a 2 Mb windows centered on the *Firre* locus and examined whether the neighboring genes were significantly dysregulated across 9 biological contexts.

Differential gene expression analysis for WT and Δ*Firre* CLPs showed that of the 12 genes within a 2 Mb window (excluding *Firre*), none were differentially expressed (Fig. 4G). Consistent with this finding, we did not observe significant changes in gene expression (2 Mb windows centered on the *Firre* locus) in seven of the eight embryonic tissues (Fig. 4G and Extended Data Fig. 9A-F). Indeed, we observed one instance of differential expression in one embryonic tissue (*Hs6st2* was slightly but significantly downregulated in the embryonic forebrain, −0.38 log2 fold change, FDR<0.05) (Extended Data Fig. 9A-F). These data demonstrate that the *Firre* locus does not exert a local effect on gene expression *in vivo*, and suggest that the *Firre* lncRNA regulates gene expression in a *trans*-based manner. Collectively, our study investigates the roles of DNA and RNA at the *Firre* locus *in vivo* and genetically defines that the *Firre* locus produces as a *trans*-acting lncRNA molecule in a hematopoetic context.

## DISCUSSION

Classic models used to study noncoding RNAs – ribosomal RNAs, small nucleolar RNAs, tRNAs, and telomerase RNA component (TERC) – have demonstrated that these RNAs species serve important cellular functions. This core of possible RNA biology has been greatly expanded by studies that have identified tens-of-thousands of lncRNAs^39,40,63^. Indeed, subsequent molecular and genetic interrogation of lncRNA loci have identified diverse molecular roles and biological phenotypes. However, lncRNA loci potentially contain multiple modes that can exert function, including DNA regulatory elements (including the promoter), the act of transcription, and the lncRNA. Therefore, attributing an RNA-based role to a lncRNA locus requires the development of multiple genetic models to determine the activities and contributions of potential regulatory modalities^20,21,59^.

In this study, we developed three genetic models in mice for the syntenically conserved lncRNA *Firre*: loss-of-function, overexpression, and rescue. Notably, we report that deletion of the *Firre* locus does not impact survival in mice, or despite escaping XCI, skew the sex ratio of progeny. We leveraged the genetic models to discriminate between DNA- and RNA-mediated effects *in vivo*. We determined that modulating *Firre* directs cell-specific defects during hematopoiesis, potentiates the innate immune response upon exposure to LPS, and can restore gene expression programs – all of which have an RNA-based functional modality. We also conclude that the *Firre* locus does not have a local *cis*-regulatory effect on gene-expression across numerous tissues. Together, by using multiple genetic and molecular approaches we identified that *Firre* produces a *trans*-acting lncRNA in a hematopoietic context. There are several important implications for these results.

First, our study indicates that *Firre* has *trans* RNA-based activity *in vivo*, and thus extends previous reports that have suggested RNA-based roles for *Firre* in cell culture models^34,47^. By using compound genetic approaches, we found that overexpression of *Firre* from a transgene in the *Firre*-deficient background was sufficient to rescue physiological and molecular phenotypes in ΔFirre CLPs *in vivo*. We speculate that early hematopoietic progenitor cells may represent a unique context to study the role(s) of *FIRRE*/*Firre*. In humans, *FIRRE* is expressed as both circular (circ-*FIRRE*) and linear forms in hematopoietic cells, and circ-*FIRRE* is abundant in all progenitor cell types except for the CLPs^64^. More studies will be needed to determine the functional differences between linear and circular isoforms of *Firre in vivo*.

Second, we observed that overexpression of *Firre* RNA in an endotoxic shock model potentiated the innate immune response *in vivo*. Our findings suggest that ectopic or high levels *Firre* lncRNA could be important for regulating the innate immune response. Consistent with our findings, modulating the levels of human and mouse *FIRRE*/*Firre* RNA in human intestinal epithelial and mouse macrophage cell lines perturbed the mRNA levels of inflammatory genes, including IL12-p40^34^. Interestingly, we identified increased levels of IL12-p40 protein in the serum from *Firre*^OE^ mice upon exposure to LPS. We speculate that *Firre* could be important in setting functional thresholds for cells. For example, *FIRRE* is not only significantly increased in certain cancers^38^, but high levels of *FIRRE* expression have been significantly associated with more aggressive disease and poor survival in patients with large B-cell lymphoma^36^.

Finally, our study suggests that *Firre* does not have a *cis*-regulatory role on gene expression *in vivo*. Upon deletion of the *Firre* locus and its promoter region we observed global changes in gene expression. Yet, we did not find changes in local gene expression (2 Mb window) at the *Firre* locus in eight embryonic tissues and in CLPs. Thus, potential DNA-regulatory elements, the lncRNA, the promoter, and the act of transcription appear to not have regulatory roles on neighboring gene expression in the nine biological contexts (8 embryonic and 1 cell type) analyzed in this study. Moreover, we observed that deletion of *Firre in vivo* does not perturb *Xist* RNA expression in eight embryonic tissues and does not affect random XCI in MEFs, consistent with a previous study using cell culture models^29^. Together, these data are notable because *cis*-acting mechanisms are speculated to be common feature at lncRNA loci^65^. While we did not find any evidence for *cis*-activity at the *Firre* locus *in vivo*, a previous study from our group found active DNA elements within the *Firre* locus using a cell-based enhancer reporter assay in 3T3 cells^22^. We speculate that these candidate DNA regulatory elements are likely to regulate the *Firre* locus rather than neighboring genes, as we did not find evidence of dysregulation in gene expression for neighboring genes when the locus was deleted *in vivo*.

In summary, we have examined the role of *Firre* in the context of hematopoiesis in order to test DNA- and RNA-mediated effects. This study does not exclude that *Firre* could be functioning elsewhere, and even by other molecular modalities. Indeed, we identified that *Firre* is abundantly expressed in a number of tissues, therefore going forward it will be important to investigate the potential role(s) of *Firre* in other biological contexts as well as in pathological disease. Our findings provide evidence that the X chromosome lncRNA locus *Firre* has a role in hematopoiesis that is mediated by a *trans*-acting RNA, and further highlights the biological importance of lncRNA-based machines *in vivo*.

## METHODS

### Mouse care and ethics statement

Mice used in these studies were maintained in a pathogen-specific free facility under the care and supervision of Harvard University’s Institutional Animal Care Committee.

### Mouse strains and genotyping

*Firre*^*floxed*^ mice were generated from 129/C57 F1 hybrid mouse embryonic cells as previously described^23^. Briefly, sequential targeting was used to insert a floxed-*neomycin*-floxed cassette in the 5’ end of the *Firre* locus between nucleotides 4790843-4790844 (mm9) and a floxed-*hygromycin*-floxed cassette was inserted into the 3’ end of the *Firre* locus between nucleotides 47990293-47990294 (mm9) (Extended Data Fig. 1A). 129/C57 F1 hybrid cells containing the *Firre*^*floxed*^ allele were injected into 129/C57 blastocysts (Harvard Genome Modification Facility). Transgenic mice were screened for the *Firre*^*floxed*^ allele by PCR genotyping. To generate a deletion of the *Firre* locus, female *Firre*^*floxed*^ mice were matted to a male *B6.C-Tg(CMV-Cre)1Cgn/J* mouse^46^ (Jackson Lab, 006054). Tail biopsies were collected from the progeny and were genotyped for WT, knockout, neomycin, hygromycin, and cre alleles (Extended Data Fig. 1B). Female mice heterozygous for the *Firre* deletion were subsequently mated to C57BL/6J mice. *Firre* WT and Δ*Firre* mice used in this study are from the F3 generation.

To generate an inducible *Firre*-overexpressing allele in mice (tg(*Firre*)), we cloned a *Firre* cDNA into a Tet-On vector (pTRE2) where the beta globin intron sequence was removed. We next used both EcoRI and NheI restriction enzymes to digest the cassette containing the tet-responsive element, CMV minimal promoter, *Firre* cDNA, and beta globin poly(A) terminator. This cassette was injected into the pronucleus of C57BL/6J zygotes (Harvard Genome Modification Facility). Male founder mice containing the tg(*Firre*) cassette were identified and individually mated to female C57BL/6J mice (Jackson Laboratory, 000664). To overexpress tg(*Firre*) F2 and F3 generation females were mated to male B6N.FVB(Cg)-Tg(CAG-rtTA3)4288Slowe/J (rtTA) mice (Jackson Laboratory, 016532) and at the plug date females were either put on a normal diet or 625 mg/kg doxycycline-containing food (Envigo, TD.01306) until experimental end points. A colony of male rtTA mice were maintained by breeding to C57BL6/J females for up to 4 generations.

Genotyping for mice was performed on tissue collected at P7. Primers used for genotyping: *Firre* wild-type allele, F-GGAGGAGTGCTGCTTACTGG, R-TCTGTGAGCCACCTGAAATG; Δ*Firre* allele, F-TCACAATGGGCTGGGTATTCTC, R- CCTGGGTCCTCTATAAAAGCAACAG; *neomycin*, F- GACCACCAAGCGAAACATC, R- CTCGTCAAGAAGGCGATAGAA; hygromycin, F-CGGAAGTGCTTGACATTGGG, R- CGTCCATCACAGTTTGCCAGTG; Cre, F- TAATCCATATTGGCAGAACG, R- ATCAATCGATGAGTTGCTTC; Sry, F- TTGTCTAGAGAGCATGGAGGGCCATGTCAA, R- CCACTCCTCTGTGACACTTTAGCCCTCCGA; tg(*Firre*) allele, F: TACCACTCCCTATCAGTGA, R: CGGCTTCATCTTCAGTCCTC; and the rtTA allele, F: AGTCACTTGTCACACAACG, R: CTCTTATGGAGATCCCTCGAC. Additional genotyping was performed by Transnetyx using real-time PCR.

### Cytokine analysis and *in vivo* endotoxin challenge

To investigate the cytokine response *in vivo*, we used two different preparations of LPS from *Escherichia coli* (*E. coli*) O111:B4. (Sigma, L2630) and Ultrapure LPS, *E. coli* O111:B4 (InvivoGen, tlrl-3pelps) and dissolved in 0.9% saline solution (Teknova, S5825). We administered either 0.9% saline or 5 mg/kg broad-acting LPS (Sigma, L2630) by i.p. injection using a 30G needle (BD insulin syringes, 328411) to mice cohorts 8 to 10 weeks old (WT, Δ*Firre, Firre*^OE^ no dox, and dox fed *Firre*^OE^). We also administered either a 0.9% saline or a 5 mg/kg dose of LPS that is TLR4-specific (InvivoGen, tlrl-3pelps) by i.p. injection in mice 5 to 10 weeks old (WT, *Firre*^OE^ no dox, and dox fed *Firre*^OE^). At 5 hours post i.p. injection, mice were euthanized and peripheral blood was collected by cardiac puncture and allowed to clot for 30 minutes at room temperature with gentle rotation. After clotting, samples were centrifuged at 1000 x g for 10 minutes at 4°C and serum was collected. Cytokine analysis was performed on serum diluted 2-fold in PBS pH7.4 (Eve Technologies, Chemokine Array 31-Plex). Measurements within the linear range of the assay are reported.

Endotoxin survival experiments were performed over two independent experiments using mice 9 to 16 weeks old: WT (mean age = 12.9 weeks), Δ*Firre* (mean age = 16 weeks), *Firre*^OE^ no dox (mean age = 13.6 weeks), and dox fed *Firre*^OE^ (mean age = 13.7 weeks). Saline control group consisting of WT, Δ*Firre*, and *Firre*^OE^ mice. 0.9% saline or 5 mg/kg LPS (InvivoGen, tlrl-3pelps) was prepared as described above and administered by i.p. injection and mice were monitored for moribund survival over 6 days. Mice were housed at a density of 3 to 5 mice per cage containing: Anderson’s Bed (The Andersons, Inc), Enviro-Dri (Shepherd Specialty Papers), compressed 2” x 2” cotton nestlet (Ancare), and a mouse hut (BioServ). The following supportive care was provided during the duration of the experiments: hydrogel, a small cup containing powdered diet mixed with water, and a heating pad (5” × 8.6” × 6”) was placed externally on the bottom of the cage (Kobayashi).

### Whole-mount *in situ* hybridization

We generated an antisense digoxigenin-labeled antisense riboprobe against *Firre* from a 428bp sequence (Extended Data Fig. 1C) corresponding to the 5’ end of the *Firre* transcript. *In situ* hybridization was performed on a minimum of three embryos per stage and/or genotype. For whole-mount staining, we fixed embryos in 4% paraformaldehyde for 18 hours at 4°C, followed by 3 washes in 1× PBS for 10 minutes at room temperature. We then dehydrated the embryos them for 5 min at room temperature in a series of graded methanol solutions (25%, 50%, 75%, methanol containing 0.85% NaCl, and 100% methanol). Embryos were then stored in 100% methanol at −20°C. We then rehydrated embryos through a graded series of 75%, 50%, 25%, methanol/ 0.85% NaCl 5 min incubations at room temperature and then washed in twice in 1× PBS with 0.1% Tween-20 (PBST). Embryos were treated with 10mg/mL proteinase K in 1× PBST for 10 minutes (E8.0, E9.5) or 30 minutes (E10.5, E11.5 and E12.5). Samples were fixed again in 4% paraformaldehyde/0.2% glutaraldehyde in PBST for 20 minutes at room temperature and washed in twice in 1× PBST. We then incubated samples in pre-hybridization solution for 1 hour at 68°C and then incubated samples in 500 ng/mL of *Firre* antisense riboprobe at 68°C for 16 hours. Post hybridization, samples were washed in stringency washes and incubated in 100 μg/mL RNaseA at 37°C for 566 1 hour. Samples were washed in 1X maleic acid buffer with 0.1% Tween-20 (MBST) and then incubated in Roche Blocking Reagent (Roche, 1096176) with 10% heat inactivated sheep serum (Sigma, S2263) for 4 hours at room temperature. We used an anti-digoxigenin antibody (Roche, 11093274910) at 1:5000 and incubated the samples for 18 hours at 4°C. Samples were washed 8 times with MBST for 15 min, 5 times in MBST for 1 hour, and then once in MBST for 16 hours at 4°C. To develop, samples were washed 3× for 5 min at room temperature with NTMT solution (100 mM NaCl, 100 mM Tris-HCl (pH 9.5), 50 mM MgCl_2_, 0.1% Tween-20, 2 mM levamisole). The *in situ* hybridization signal was developed by adding BM Purple (Roche, 11442074001) for 4, 6, 8, and 12 hours. After the colorimetric development, samples were fixed in 4% paraformaldehyde and cleared through a graded series of glycerol/1× PBS and stored in 80% glycerol. Embryos were imaged on a Leica M216FA stereomicroscope (Leica Microsystems) equipped with a DFC300 FX digital imaging camera.

### RNA-seq in embryonic tissues preparation and analysis

For WT and Δ*Firre* RNA-seq in embryonic tissues we dissected tissues (forebrain, midbrain, heart, lung, liver, forelimb, hindlimb, and presomitic mesoderm) from E11.5 embryos (44-48 somites) that were collected from matings between either male WT and female *Firre*^+/-^ or male Δ*Firre* and female *Firre*^+/-^ mice. Tissues were immediately homogenized in Trizol (Invitrogen) and total RNA was isolated using RNeasy mini columns (Qiagen) on a QIAcube (Qiagen). Samples were genotyped for the WT, Δ*Firre*, and sex alleles (Extended Data Fig. 1A,B). For each tissue, we generated the following libraries: WT male (*n*=3), WT female (*n*=3), Δ*Firre* male (*n*=3), and Δ*Firre* female (*n*=3). Poly(A)+ RNA-seq libraries were constructed using TruSeq RNA Sample Preparation Kit v2 (Illumina). The libraries were prepared using 500ng of total RNA as input, with the exception of the lung (200ng) and the presomitic mesoderm (80ng), and with a 10-cycle PCR enrichment to minimize PCR artifacts. The indexed libraries were pooled in groups of six, with each pool containing a mix of WT and Δ*Firre* samples. Pooled libraries were sequenced on an Illumina HiSeq 2500 in rapid-run mode with paired-end reads.

Reads were mapped to the mm10 mouse reference genome using TopHat v2.1.1 with the flags: “--no-coverage-search --GTF gencode.vM9.annotation.gtf” where this GTF is the Gencode vM9 reference gene annotation available at gencodegenes.org. Cufflinks v2.2.1 was used to quantify gene expression and assess the statistical significance of differences between conditions. Cuffdiff was used to independently compare the WT and Δ*Firre* samples and from each tissue and sex, and genes with FDR<0.05 were deemed significant (Extended Data Tables 2-9).

### RNA-seq in CLPs preparation and analysis

We isolated CLPs (Lin-Sca-1^lo^cKit^lo^IL7Rα+) by fluorescence activated cell sorting (FACS) from mice 27 to 32 weeks old: WT (*n*=4, mean age = 31 weeks), Δ*Firre* (*n*=4, mean age = 30.4 weeks), and dox fed *Firre*^rescue^ (*n*=4, mean age = 29.3 weeks. CLPs were directly sorted into TRIzol. RNA was isolated using RNeasy micro columns (Qiagen) on a QIAcube (Qiagen) and we quantification the concentration and determined the RNA integrity using a BioAnalyzer (Agilent). Poly(A)+ RNA-seq libraries were constructed using CATS RNA-seq kit v2 (Diagenode, C05010041). Pooled libraries were sequenced on an Illumina HiSeq 2500 in rapid-run mode with paired-end reads.

The adapter-trimmed reads were mapped to the mm10 mouse reference genome using TopHat v2.1.1 with the flags: “--no-coverage-search --GTF gencode.vM9.annotation.gtf” (Gencode vM9 reference gene annotation) FeatureCounts and R-package, DESeq2, were used to quantify gene expression and assess the statistical significance of differences between conditions^66,67^ and the p-value of comparisons were empirically calculated by using fdrtools^68^. Genes with an FDR<0.1 were deemed significant in a comparison between wildtype and Δ*Firre* (Extended Data Table 10) and genes with an FDR<0.1 in the Δ*Firre* and *Firre*^rescue^ comparison were deemed significant (Extended Data Table 11).

### qRT-PCR

Embryonic tissues were homogenized in Trizol (Invitrogen) and total RNA was isolated using RNeasy mini columns (Qiagen) on a QIAcube (Qiagen). 300ng of total RNA was used as input to synthesize cDNA (SuperScript IV VILO Master Mix, Invitrogen, 11756050). Primers used qRT-PCR experiments: F_b-act: GCTGTATTCCCCTCCATCGTG, R_b-act: CACGGTTGGCCTTAGGGTTCAG; F_Firre: AAATCCGAGGACAGTCGAGC, R_Firre: CCGTGGCTGGTGACTTTTTG. Experiments were performed on a Viia7 (Applied Biosciences). qRT-PCR data was analyzed by the ΔΔCt method.

### Distribution of *Firre* expression across wild-type tissues

For each of the eight WT embryonic tissues, FPKM estimates of all protein coding or noncoding genes were aggregated and filtered for expression > 1FPKM. Density plots were generated using ggplot2 (geom_density()). The distributions of wildtype FPKM estimates for these transcriptional types are indicated by black line for protein-coding and gray line for non-coding expression.

### Flow Cytometry analysis

Age and sex-matched adult mice were used in all flow cytometry experiments. Zombie Aqua Fixable Viability Kit (Biolegend, 423101) was used as a live-dead stain. CountBright Absolute Counting Beads (Invitrogen, C36950) were added to bone marrow and thymi samples in order to enumerate cell populations.

For cell analysis, peripheral blood was collected by cardiac puncture. The following antibodies were added (1:100) to each sample and incubated for 30 minutes at room temperature Alexa Fluor 700 anti-mouse CD8a (Biolegend, 100730), PE/Dazzle-594 anti-mouse CD4 (Biolegend, 100456), APC anti-mouse CD19 (Biolegend, 115512), Alexa Fluor 488 anti-mouse NK-1.1 (Biolegend, 108718), PE anti-mouse CD3 (Biolegend, 100205), PE/Cy7 anti-mouse/human CD44 (Biolegend, 103030), eFluor 450 anti-Mouse CD62L (L-Selectin, eBiosciences, 48-0621-82), and TruStain FcX (anti-mouse CD16/32) antibody (1:50) (Biolegend, 101319). Red blood cells were then lysed for 15 minutes at room temperature using BD FACS Lysing Solution (BD, 349202). Cells were washed twice in 1× PBS with 1% BSA and then resuspended in 1% paraformaldehyde or 1× PBS with 0.2% BSA.

Thymi were collected and homogenized in ice cold PBS over a 40 micron filter. The cells were incubated with the following antibodies (1:100) for 30 minutes at room temperature: Alexa Fluor 488 anti-mouse CD25 (Biolegend, 102017), PE/Cy7 anti-mouse/human CD44 (Biolegend, 103030), PE anti-mouse TCR β chain (Biolegend, 109208), APC anti-mouse/human CD45R/B220 (Biolegend, 103212), eFluor 450 anti-Mouse CD69 (eBiosciences, 48-069182), Alexa Fluor 700 anti-mouse CD8a (Biolegend, 100730), and PE/Dazzle-594 anti-mouse CD4 (Biolegend, 100456). Cells were washed twice in 1x PBS with 1% BSA and then resuspended in 1% paraformaldehyde.

Bone marrow was collected from both femurs and tibias (four bones total per mouse) by removing the end caps and flushing with DMEM (Gibco, 11995-073) containing 5% FBS (Gibco, 26140079) and 10mM EDTA. Cells were then pelleted, re-suspended and passed through a 70 micron filter. The resulting single cell suspension was then incubated with the following antibodies (1:100) for 60 minutes on ice: Alexa Fluor 700 anti-Mouse CD16/CD32 (eBiosciences, 65016182), PE/Cy7 anti-mouse CD127 (IL-7Rα) (Biolegend, 135014), Alexa Fluor 488 anti-mouse CD117 (c-Kit) (Biolegend, 105816), PE/Dazzle-594 anti-mouse Ly-6A/E (Sca-1) (Biolegend, 108138), Pacific Blue anti-mouse Lineage Cocktail (Biolegend, 135306), PE anti-mouse CD135 (Biolegend, 135306), and APC anti-mouse CD34 (Biolegend, 128612). Red blood cells were then lysed for 15 minutes at room temperature using BD FACS Lysing Solution (BD, 349202) or BD Pharm Lyse (BD, 555899). Cells were washed twice in 1× PBS with 1% BSA and then resuspended in 1% paraformaldehyde or 1× PBS with 0.2% BSA.

Flow cytometry was performed on a LSR-II (BD) and the gating was performed using FlowJo software (Treestar) using the following criteria (applied to live singlets): CD4 T cells (CD3+CD4+CD8-CD19-; CD8 T cells (CD3+CD8+CD4-CD19-); NK cells (NK1.1+B220-CD3-); B cells (CD19+CD3-); double negative (DN) (B220-CD4-CD8-CD25^var^CD44^var^); double positive (DP) (CD4+CD8+B220-); single positive (SP) (CD8, CD8+CD4-B220-); single positive (SP) (CD4, CD4+CD8-B220-); hematopoietic stem cells (HSC) (LSK [Lin-, Sca-1+, c-Kit+)-CD34+-CD135-); multipotent progenitors (MPP) (LSK-CD34+CD135+); common lymphoid progenitors (CLP) (Lin-Sca-1^lo^c-Kit^lo^IL7Rα+); and common myeloid progenitors (CMP) (CD34+CD16/32-). Negative gates were set using fluorescence-minus-one controls (FMO).

### Competitive HSC transplant assay

Bone marrow from age- and sex-matched mice was collected and pooled with like genotypes (as described in flow cytometry analysis section) from mice that were 8 to 9 weeks in age: PepBoy/CD45.1 (*n*=3 females per experiment; mean age = 9 weeks) (Jackson Laboratory, 002014), *Firre* WT/CD45.2 (*n*=3 females per experiment mean age = 8.9 weeks), and Δ*Firre*/CD45.2 (*n*=3 females per experiment; mean age = 8.6 weeks). Bone marrow was lineage depleted according to the manufacture protocol (MiltinyiBiotec, 130-042-401), and cell marker surface staining was performed (as described for bone marrow in flow cytometry analysis section). Red blood cells were then lysed for 15 minutes at room temperature using BD Pharm Lyse (BD, 555899). Cells were washed twice in 1× HBSS with 5% FBS and 2 mM EDTA. We then double sorted lineage depleted cells for an HSC-enriched population (Lin^-^ Sca-1^+^c-Kit^+^CD34^+-^CD135^-^) into 2% FBS in HBSS using fluorescent activated cell sorting (FACS) (BD Aria). Recipient mice, PepBoy/CD45.1 (*n*=10 males per experiment; mean age = 8.6 weeks), were lethally irradiated using a split 9.5γ split dose (3 hours apart). *Firre* WT and Δ*Firre* HSCs were mixed at a 1:1 ratio to PepBoy/CD45.1 HSCs and 100 ul containing 4,000 cells were transplanted by retro-orbital injection using 30-gauge insulin syringes (BD, 328411) into the lethally irradiated recipients. 48 hours post-transplant, 100,000 helper marrow cells from male PepBoy/CD45.1 were transplanted by retro-orbital injection into each experimental PepBoy/CD45.1 male recipient mouse.

### MEF preparations and culture

We generated *Firre* WT, *Firre* knockout, and *Firre*^rescue^ MEFs at E13.5 from intercrosses between male *Firre*^-/y^ with female *Firre*^+/-^ and male *Firre*^rescue^ with female *Firre*^-/-^. Individual embryos were dissected into 1× phosphate-buffered saline (PBS) and were eviscerated, and the head, forelimbs, and hindlimbs were removed. Embryo carcasses were placed into individual 6 cm^2^ tissue culture plates containing 1 mL of pre-warmed 37°C TrypLE (Thermo Fisher, 12604013) and were incubated at 37°C for 20 min. Embryos were dissociated by gently pipetting using a P1000 tip and MEF media was added. Cells were cultured for 5 to 7 days and cryostocks of individual lines were generated. Subsequent experiments were performed from thaws from the cryostocks up to passage 3. MEFs were genotyped for *Firre* WT, knockout, rtTA, tg(*Firre*), and sry alleles. MEF culture media: 1× Dulbecco’s modified Eagle’s medium (DMEM) (Invitrogen 11965-118), 10% fetal bovine serum (Gibco, 10082139), L-glutamine (Thermo Fisher, 25030081), and penicillin/streptomycin (Thermo Fisher, 15140122).

### *Firre* RNA FISH

*Firre* WT, Δ*Firre*, and *Firre*^rescue^ MEFs were plated at a density of 50,000 cells per well onto round glass cover slips in a 24-well plate. *Firre*^rescue^ MEFs were cultured with either 2 ug/mL dox (Sigma, D9891) or vehicle (ddH_2_O) for 24 hours. Replicate wells were processed for either RNA FISH or to isolate RNA for *Firre* induction analysis by qRT-PCR. RNA FISH using oligo probes was performed as previously described^69^. Briefly, *Firre* oligo probes were designed using Primer3 (http://frodo.wi.mit.edu/primer3/) and synthesized by Integrated DNA Technologies. After Amine-ddUTP (Kerafast) was added to 2 pmol of pooled oligos by terminal transferase (New England Biolabs), oligos were labeled withAlexa647 NHS-ester (Life Technologies) in 0.1 M sodium borate. Cells grown on glass coverslips were rinsed in PBS and fixed in 4% paraformaldehyde. After permeabilization in 0.5% Triton X-100 at room temperature, cells were washed in PBS and dehydrated in a series of increasing ethanol concentrations. 6 labeled oligo probes were added to hybridization buffer containing 25% formamide, 2× SSC, 10% dextran sulfate, and 1 mg/mL yeast tRNA. RNA FISH was performed in a humidified chamber at 42°C for 4 hours. After being washed three times in 2X SSC, cells were mounted for wide-field fluorescent imaging or dehydrated for STORM imaging. Nuclei were counter-stained with Hoechst 33342 (Life Technologies). The following pooled oligos against *Firre* were used: (1) AGCAGCAAATCCCAGGGGCC, (2) TTCCTCATTCCCCTTCTCCTGG, (3) CCCATCTGGGTCCAGCAGCA, (4) ATCAGCTGTGAGTGCCTTGC, (5) TCCAGTGCTTGCTCCTGATG, (6) GCCATGGTCAAGTCCTGCAT

### *Firre* DNA/RNA and *Xist* RNA co-FISH

Primary MEF cells were trypsinzed and cytospun to glass slides. After brief air drying, cells were incubated in PBS for 1min, CSK/0.5% Triton X-100 for 2 min on ice, and CSK for 2 min on ice. Cells were fixed in 4% formaldehyde in PBS for 10 min at RT and washed twice in PBS. After dehydrated through series of EtOH, cells were subject to hybridization at 37°C O/N with denatured digoxigenin-labeled *Xist* probe (50% formamide, 2× SSC, 10% dextran sulfate, 0.1 mg/mL CoT1 DNA). Cells were washed in 50% formamide, 2X SSC at 37 ° C and in 2× SSC at RT, three times each. RNA FISH signal was detected by incubating FITC-labeled anti-digoxygenin antibody (Roche) in 4× SSC, 0.1% Tween-20 at 37 °C for 1 hour and followed by washing in 4× SSC, 0.1% Tween-20 at 37 °C three times. Cells were fixed again in 4% formaldehyde in PBS for 10 min and washed twice in PBS. Cellular RNAs were removed by RNase A (Life Technology) in PBS at 37°C. After dehydrated through series of EtOH, cells were sealed in hybridization buffer (50% formamide, 2X SSC, 10% dextran sulfate, 0.1 mg/mL CoT-1 DNA) containing Cy3-labeled *Firre* probe (Fosmid WI-755K22). Chromosomal DNA and probes were denaturated at 80°C for 15 min and allowed to renature by cooling down to 37° C O/N. Cells were washed in 50% formamide, 2X SSC at 37°C and in 2X SSC at RT, three times each. Nuclei were counter-stained with Hoechst 33342 (Life Technology). Imaging was performed on Nikon 90i microscope equipped with a 60X/1.4 N.A. VC objective lens, Orca ER camera (Hamamatsu) and Volocity software (Perkin Elmer). All probes were prepared by nick translation using DNA polymerase I (New England Biolab), DNase I (Promega), and Digoxigenin-dUTP (Roche), or Cy3-dUTP (Enzo Life Sciences)

### Skeletal preparations

WT and Δ*Firre* E18.5 embryos were dissected and eviscerated. Samples were fixed in 100% ethanol for 24 hours at room temperature. Embryos were then placed in 100% acetone for 24 hours at room temperature and then incubated in staining solution (0.3% alcian blue 8GS (Sigma) and 0.1% Alizarin Red S (Sigma) in 70% ethanol containing 5% acetic acid) for three days at 37°C. Samples were then rinsed with distilled water and then placed in 1% potassium hydroxide at room temperature for 24 hours. Samples were then cleared in a series of 1% potassium hydroxide / 20%, 50%, and 80% glycerol.

## ACKNOWLEDGEMENTS

We thank Dr. Martin Sauvageau for providing a *Firre* clone to generate a riboprobe; Dr. Diana Sanchez for assistance in the mouse facility; Dr. Susan Carpenter, Dr. Kate Pritchett-Corning, and Elektra Robinson for discussions on the LPS study; Joyce LaVecchio and Silvia Ionescu in the HSCRB flow cytometry core for FACS assistance; the Harvard Bauer Core for sequencing; Dr. Marta Mele for initial optimization for RNA-seq analysis; and Dr. Laurie Chen and Dr. Lin Wu at the Harvard Genome Modification Facility. This research was supported by the National Institutes of Health (NIH) General Medical Sciences postdoctoral fellowship award 1F32GM122335-01A1 (to J.P.L) and support from NIH National Heart, Lung, and Blood Institute T32HL007893; NIH postdoctoral fellowship F32AG050395 (to J.M.G.); R.A.F is supported by the Howard Hughes Medical Institute; NIH RO1 AG048917 and the Dean’s Initiative Award Program for Innovation Grants in the Basic and Social Sciences (to A.J.W). the Institute of Mental Health grant R01MH102416-03 and the NIH Institute of General Medical Sciences grant P01GM099117 (to J.L.R).

## AUTHOR CONTRIBUTIONS

Study conceptualization and design: J.P.L, J.C.L, and J.L.R; *Firre* ES cell targeting: A.W., J.H., R.A.F; Transgenic mice generation and mouse husbandry, N.C. and J.P.L; Immunophenotyping experiments: J.P.L, J.C.L, and J.M.G.; Competitive chimera design and analysis: J.M.G, J.P.L, A.J.W; Endotoxic shock experiments: J.P.L, N.C, and C.G; RNA-sequencing design and analysis: T.H, J.P.L, C.G, W.M, A.G; RNA FISH for *Firre*: H.S and J.T.L; Funding and supervision: A.J.W and J.L.R; Writing manuscript J.P.L, J.C.L, and J.L.R with input from all of the authors.

## EXTENDED DATA FIGURES

**Extended Data Figure 1:**
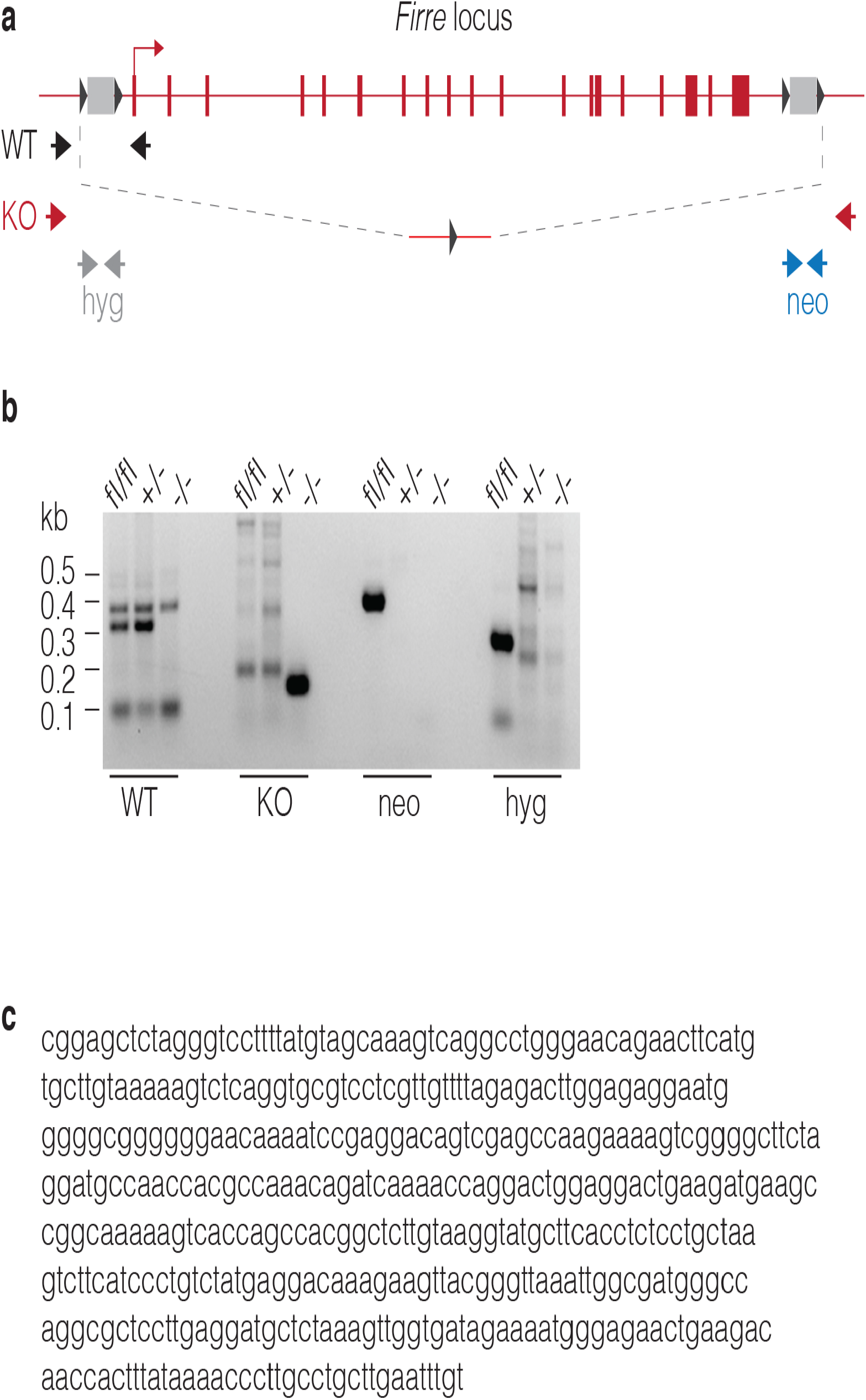
Schematization of the targeted *Firre* locus and genotyping. **(A)** Targeted *Firre* locus as described in^23^ shown in reverse orientation. Targeting cassettes containing hygromycin and neomycin cassettes shown as light gray rectangles and the loxP sites shown as dark gray triangles. Cre-mediated recombined allele shown below as a red line with a single loxP site. Arrows indicate genotyping primers used to amplify alleles for: *Firre* WT, black; knockout allele (KO), red; *hygromycin* (*hyg*), light gray; and *neomycin* (*neo*), blue. **(B)** Genotyping gel for: *Firre*^*floxed*^ (fl/fl); *Firre* heterozygous (+/-); and *Firre* knockout (-/-) mice. Primers used to amplify different alleles indicated below the gel. **(C)** DNA sequence used to generate a *Firre* riboprobe.

**Extended Data Figure 2.**
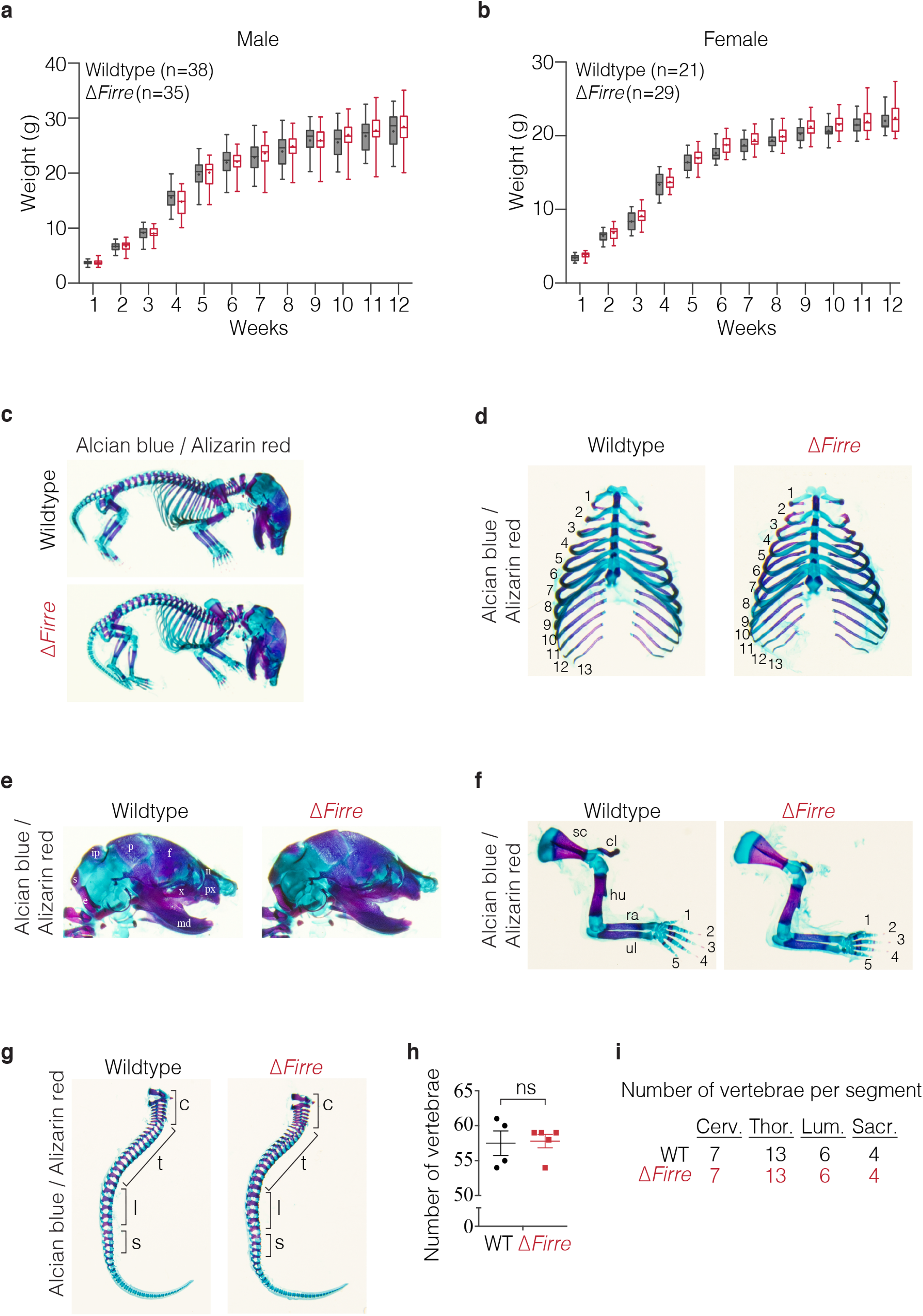
Weight measurements and skeletal analysis of Δ*Firre* mice. **(A)** Body weight measurements in male WT (*n*=38) and Δ*Firre* (*n*=35) and **(B)** female WT (*n*=21) and Δ*Firre* (*n*=29) mice over 12 weeks are not significantly different. Data shown as a box and whisker plot with the minimum and maximum, the significance was determined using a two-tailed t-test, WT shown in dark gray and Δ*Firre* shown in red. **(C-G)** Skeletal preparations of E18.5 WT (*n*=8) and Δ*Firre* (*n*=7) mice stained with alcian blue and alizarin red show that Δ*Firre* mice have normal skeletal development **(D)** Rib cages from E18.5 wild-type (*n*=8) Δ*Firre* (*n*=7) showing that Δ*Firre* embryos have a normal number of ribs. **(E)** Skulls from E18.5 WT (*n*=8) and Δ*Firre* (*n*=7) embryos show normal morphology. Abbreviations used: n, nasal; f, frontal bone; p, parietal; ip, interparietal; s, supraoccipital; e, exoccipital; md, mandible; and x, maxillary. **(F)** Limb patterning and ossification appears normal in WT (*n*=8) and Δ*Firre* mice (*n*=7). Abbreviations used: sc, scapula; cl, clavicle; hu, humerus; ra, radius; and ul, ulna. **(G)** Vertebrae patterning and ossification appears normal in WT (*n*=8) and Δ*Firre* (*n*=7) embryos. **(H)** The total number of vertebrae in E18.5 WT (*n*=4) and Δ*Firre* (*n*=5) embryos do not significantly differ (two-tailed unpaired t-test, P=0.876). Error bars indicate the SEM. **(I)** The number of vertebrae per: c, cervical; t, thoracic; l, lumbar, and s, sacral segments in E18.5 Δ*Firre* (*n*=5) embryos is the same as found in WT (*n*=4)

**Extended Data Figure 3.**
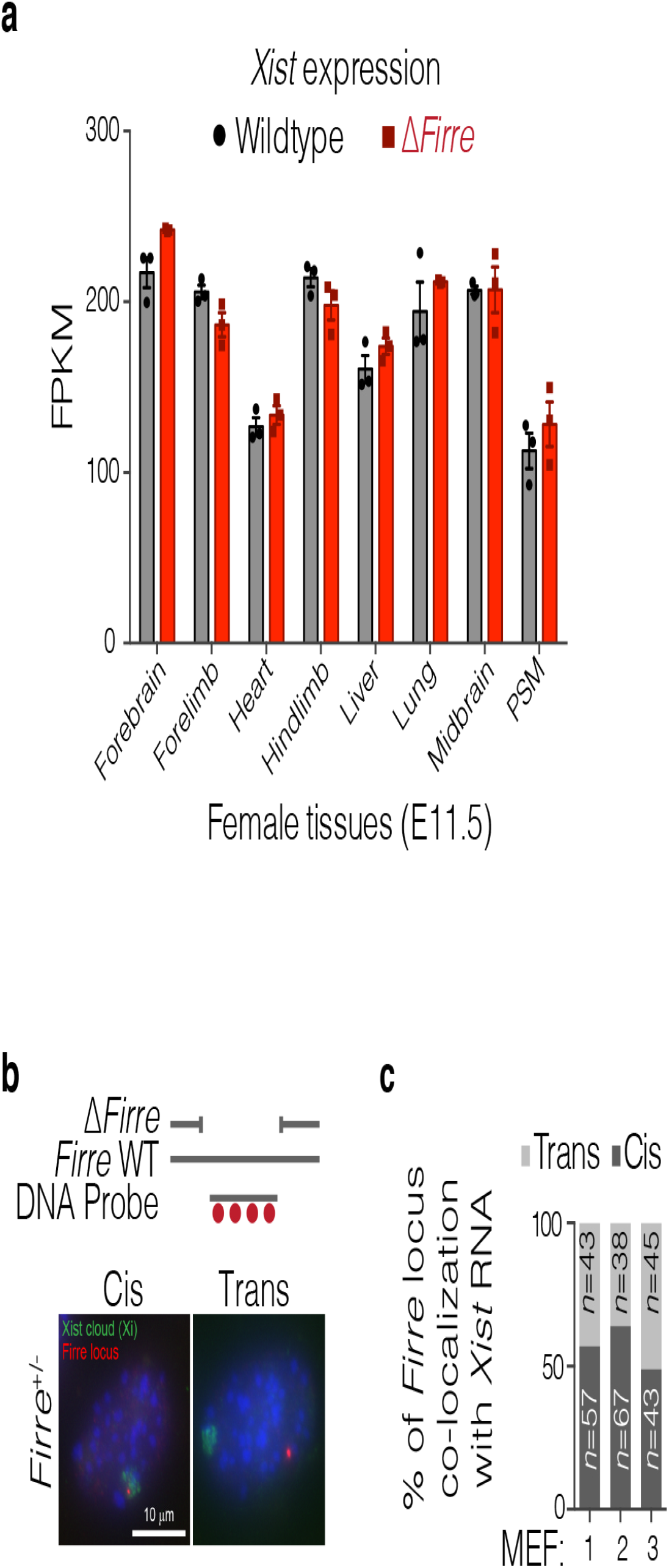
Deletion of *Firre* does not impact X chromosome inactivation or change expression of *Xist* RNA. **(A)** *Xist* RNA expression (FPKM) in eight female tissues from RNA-seq in WT (*n*=3) and Δ*Firre* (*n*=3) at E11.5. Data are shown as mean ± SEM. **(B**,**C)** Co-DNA/RNA FISH in female *Firre*^+/-^ MEFs. DNA FISH for the WT *Firre* locus shown in red and *Xist* RNA shown in green. Quantification of localization of *Xist* RNA with the WT *Firre* locus from independent *Firre*^*+/-*^ MEFs. Cis indicates a co-localization between the WT *Firre* DNA locus and *Xist* RNA and trans indicates *Xist* RNA did not co-localize with the WT *Firre* DNA locus.

**Extended Data Figure 4.**
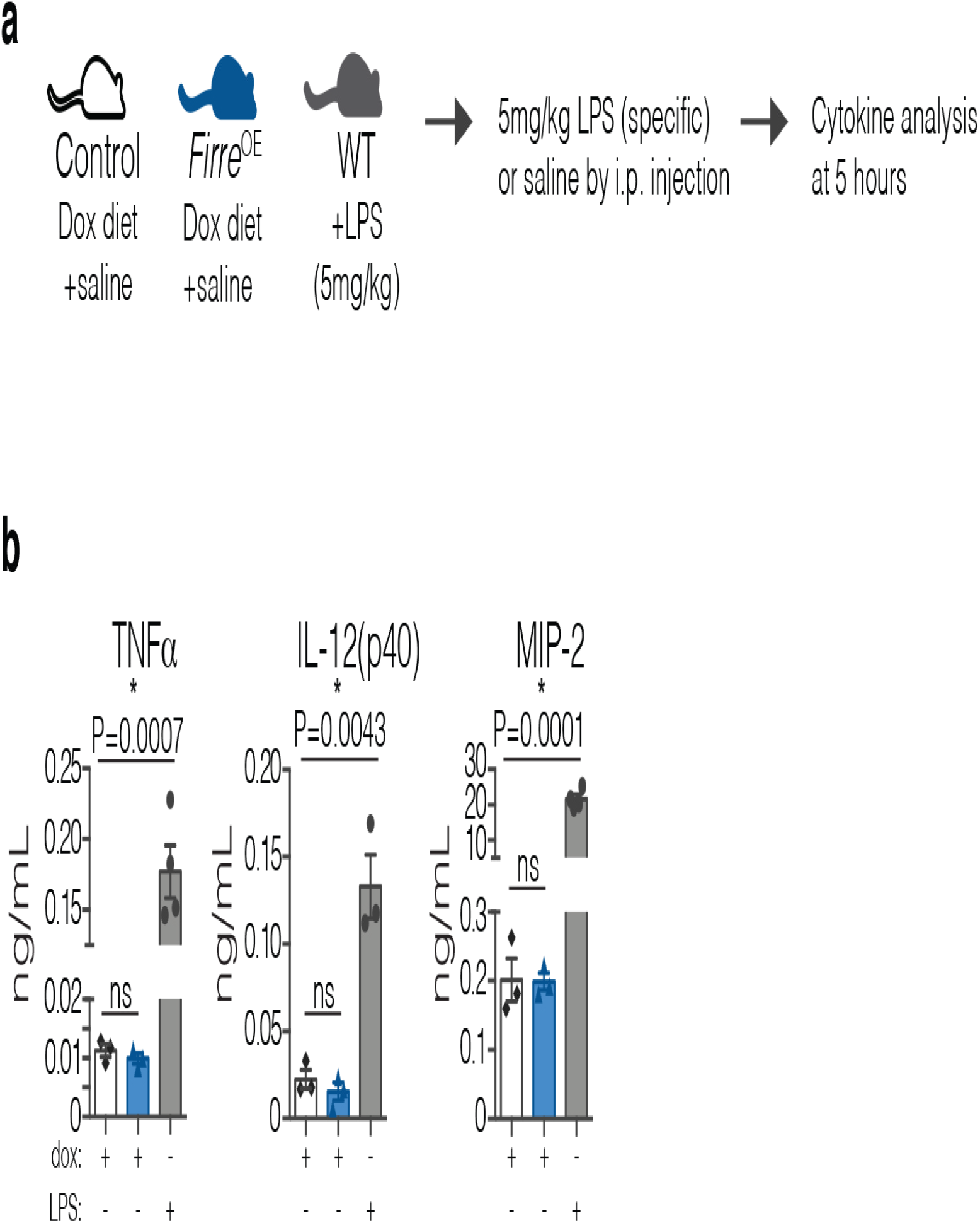
*Firre*^OE^ mice unchallenged do not have increased levels in serum cytokines. **(A)** Experimental schematic for cytokine measurements in 5-7 weeks old mice injected with either saline or LPS. **(B)** Cytokine measurements in serum at 5 hours post saline or LPS injection from control saline injected mice (WT or tg(*Firre*) fed a dox diet, *n*=3, black diamonds), *Firre*^OE^ saline injected mice fed a dox diet (*n*=3, blue triangles), and WT mice fed a normal diet injected with 5 mg/kg LPS (specific-activity) (*n*=3 to 4, gray circles). Data are shown as mean ± SEM and statistical significance determined using a two-tailed unpaired t-test.

**Extended Data Figure 5.**
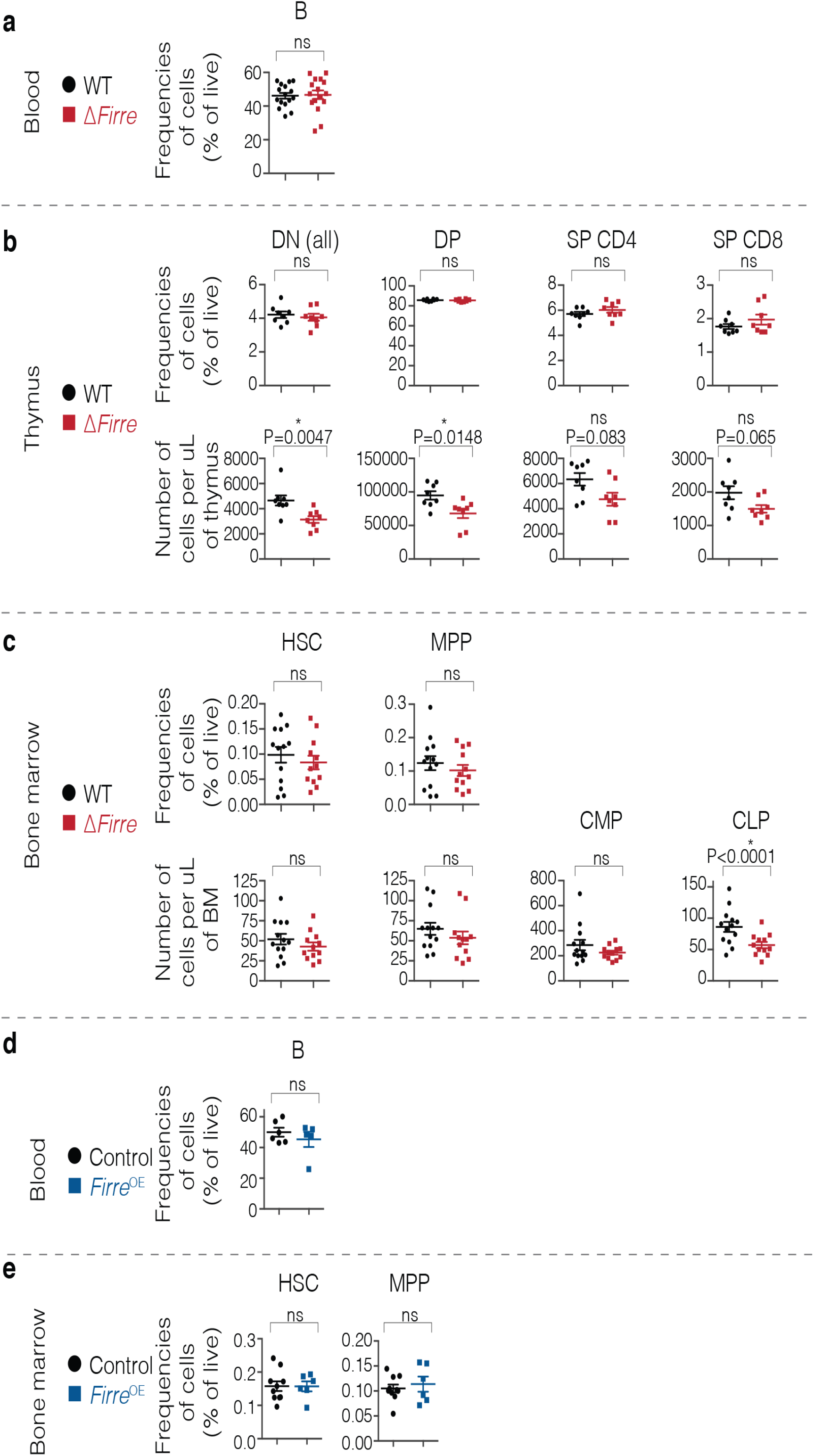
Immunophenotyping in WT, Δ*Firre*, and *Firre*^OE^ mice. **(A)** Frequency of B cells in the peripheral blood shown as percent (%) of live cells from WT and Δ*Firre* mice. Three representative experiments combined (seven independent experiments). **(B)** Frequencies of double negative (DN) (DN1, DN2, DN3, DN4), double positive (DP), single positive (SP) CD4, and SP CD8 cells in thymuses shown as percent of live cells from WT and Δ*Firre* mice. Enumeration of cells shown below as cells / uL of thymus. A representative experiment shown (three independent experiments). **(C)** Frequencies of HSC and MPP cell populations from total bone marrow (BM) shown as percent of live cells from WT and Δ*Firre* mice. Enumeration of cells shown below as cells / uL of bone marrow. Two representative experiments combined (three independent experiments). **(D)** Frequency of B cells in the peripheral blood shown as percent of live cells from control (tg(*Firre*), WT, or rtTA with dox) and dox-treated *Firre*^OE^ mice. One representative experiment shown (three independent experiments). **(E)** Frequencies of HSC and MPP cells from total BM shown as percent of live cells form control (tg(*Firre*), WT, or rtTA with dox) and dox-treated *Firre*^OE^ mice (two independent experiments). All data shown as mean ± SEM and statistical significance determined using a two-tailed Mann Whitney-U test.

**Extended Data Figure 6.**
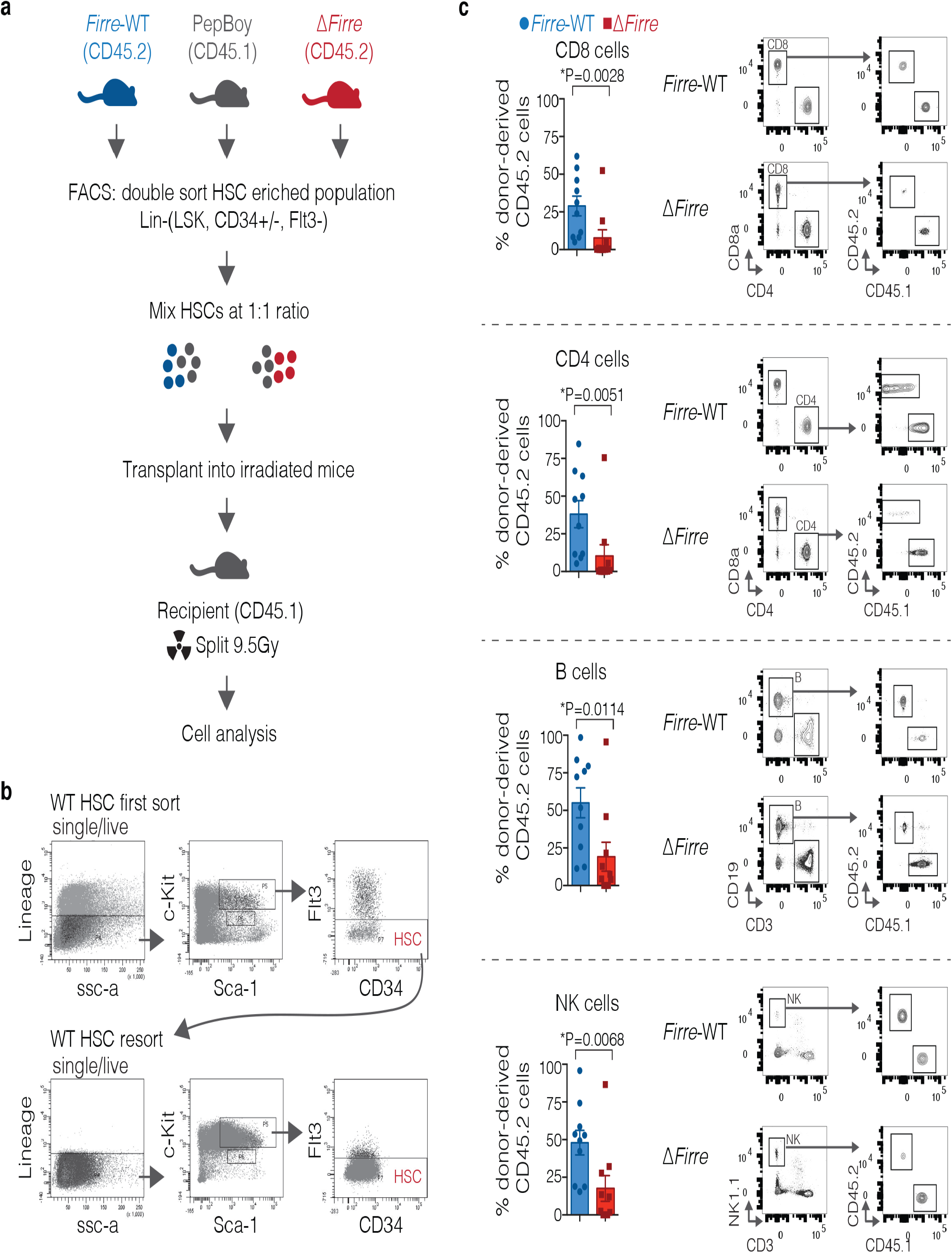
Δ*Firre* HSC populations are less competitive at repopulating the blood *in vivo*. **(A)** Schematic of competitive chimera HSC transplant experiment. HSC enriched population from age- and sex-matched *Firre* WT/CD45.2 (blue) or Δ*Firre*/CD45.2 (red) combined with PepBoy/CD45.1 (gray) at a 1:1 ratio and transplanted into lethally irradiated PepBoy/CD45.1 recipient male mice. **(B)** Representative flow cytometry plots from WT showing the FACS strategy used for isolating an HSC-enriched population for transplant from lineage depleted total bone marrow. **(C)** Frequencies of donor-derived CD45.2 CD4, CD8, NK, and B cells at 23 weeks post competitive chimera transplant for *Firre* WT/CD45.2 with PepBoy/CD45.1 (*n*=10), and Δ*Firre*/CD45.2 with PepBoy/CD45.1 (*n*=10) (two independent experiments shown). Data are shown as mean ± SEM and significance determined by a two-tailed Mann-Whitney U test.

**Extended Data Figure 7.**
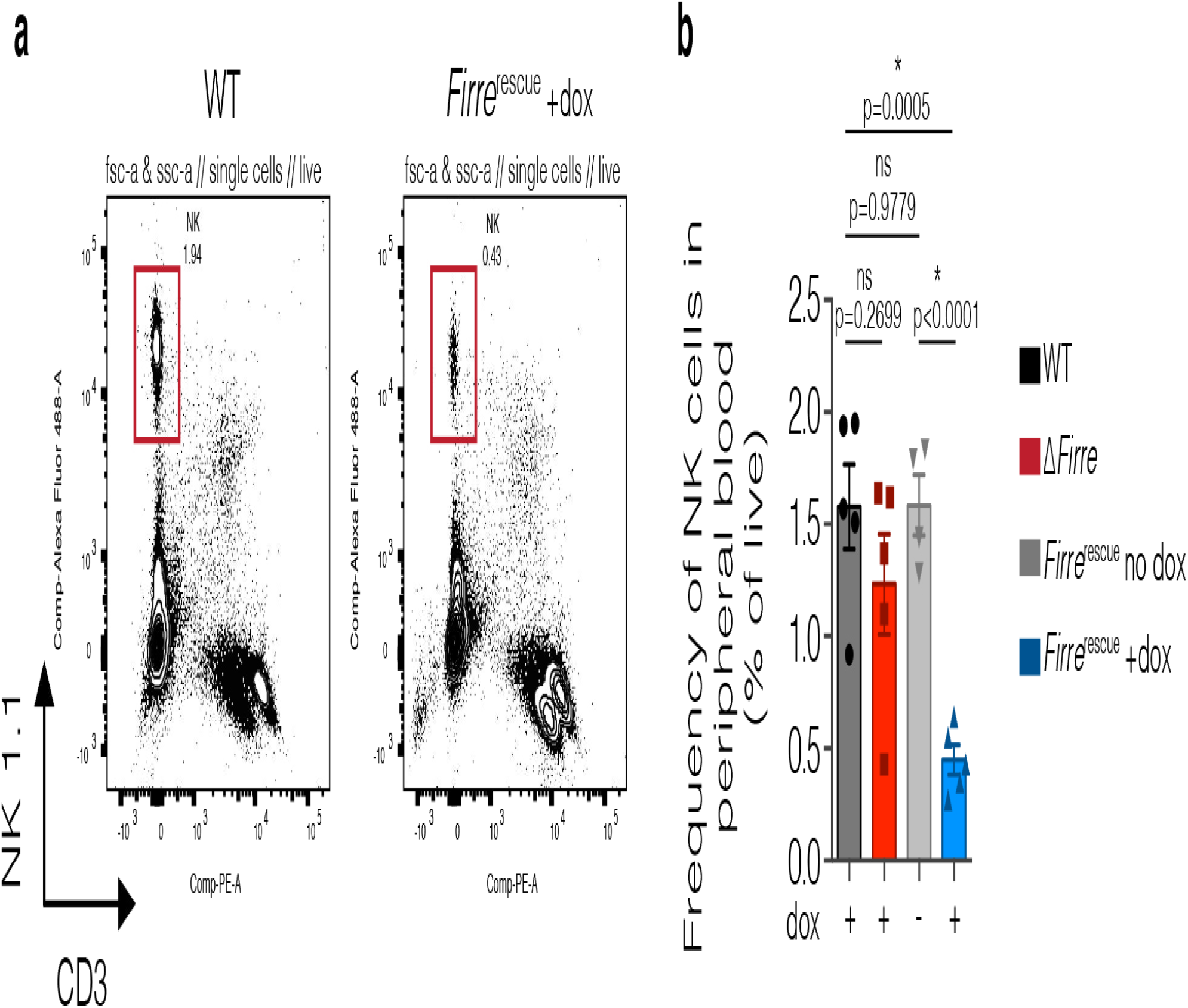
*Firre*^rescue^ mice overexpressing *Firre* RNA have a decrease in the frequency of NK cells in the peripheral blood. **(A)** Representative flow cytometry plots of NK cells in WT and dox-treated *Firre*^rescue^ mice. **(B)** Frequency of NK cells shown as percent live cells in the peripheral blood from female mice 24 to 33 weeks old: dox-treated WT (*n*=5), dox-treated Δ*Firre* (*n*=5), no dox *Firre*^rescue^ (n=4), and dox-treated *Firre*^rescue^ (*n*=5). Data are shown as mean ± SEM, two independent experiments, and significance determined by using a two-tailed unpaired t-test.

**Extended Data Figure 8.**
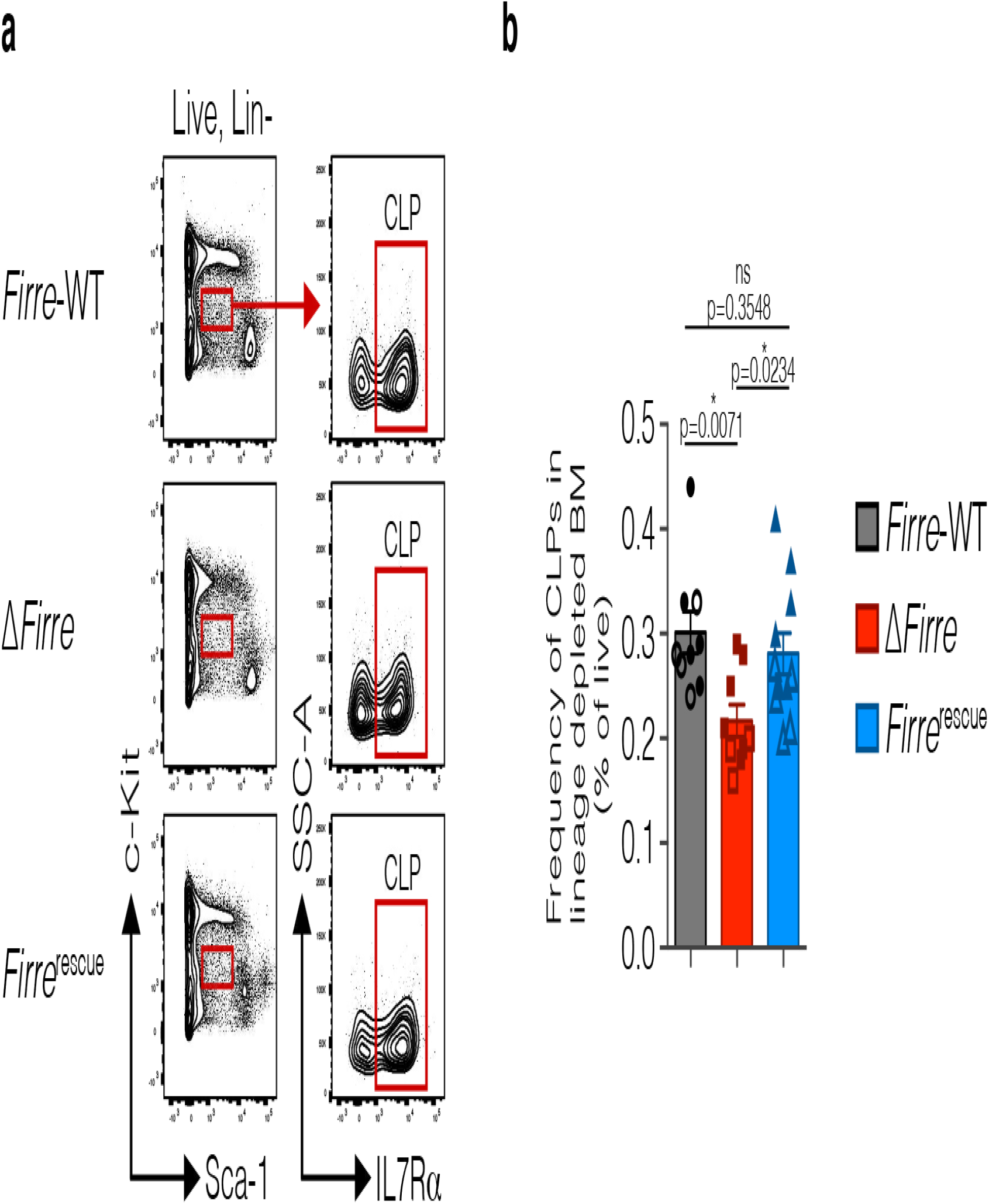
Overexpression of *Firre* RNA in *Firre*^rescue^ mice restores CLP frequency in lineage-depleted bone marrow. **(A)** Representative gating strategy for identifying CLPs in total and lineage depleted bone marrow (BM) from WT, Δ*Firre*, and dox-treated *Firre*^rescue^ mice. **(B)** Frequency of CLPs shown as percent of live cells in lineage depleted bone marrow over three experiments from male (7 to 10 weeks old, solid object) and female (19 to 24 weeks old, outlined object) mice: WT (*n*=9); Δ*Firre* (*n*=9), and dox-treated *Firre*^rescue^ (*n*=11). Data are plotted as the mean ± SEM and significance determined by a two-tailed Mann-Whitney U test.

**Extended Data Figure 9.**
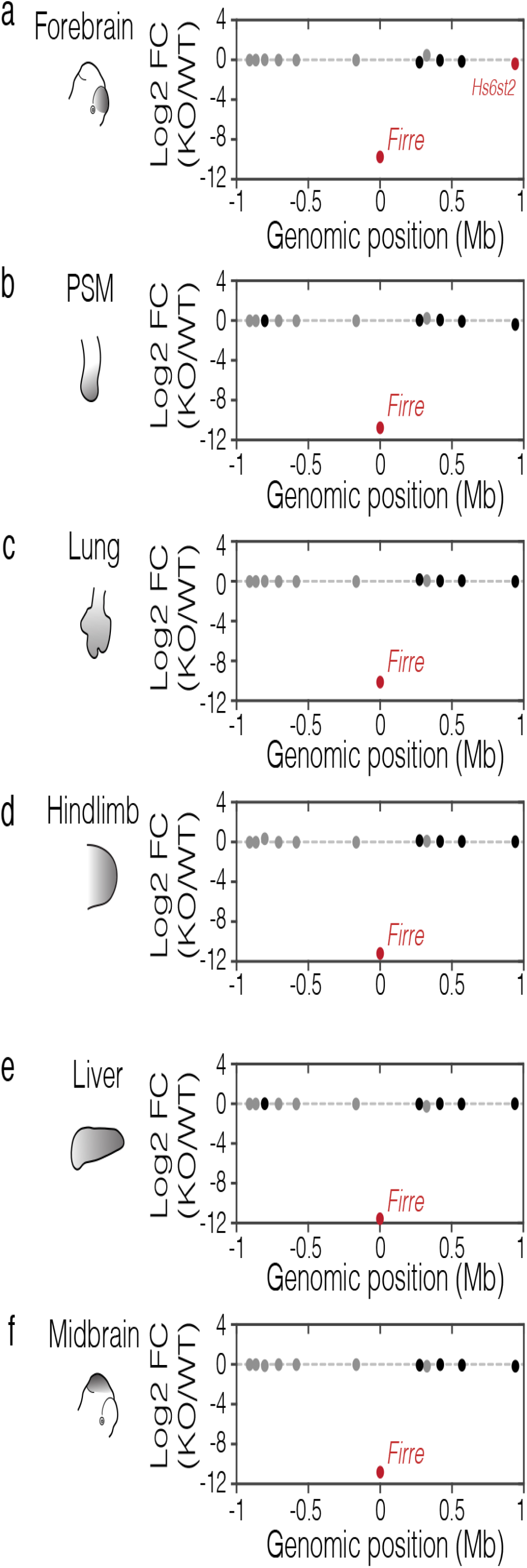
*Firre* does not regulate the expression of neighboring genes. **(A-F)** *Firre* locus region (2 Mb) showing log2 fold change (log2 FC) gene expression differences (RNA-seq) between Δ*Firre* and WT E11.5 tissues (forebrain, pre-somitic mesoderm (PSM), lung, hindlimb, liver, and midbrain). *Firre* is shown in red, significantly dysregulated genes are shown in red, genes with less than 1 FPKM expression are shown in gray, and genes that are not significantly changed are shown in black.

## EXTENDED DATA TABLES

**Extended Data Table 1.**
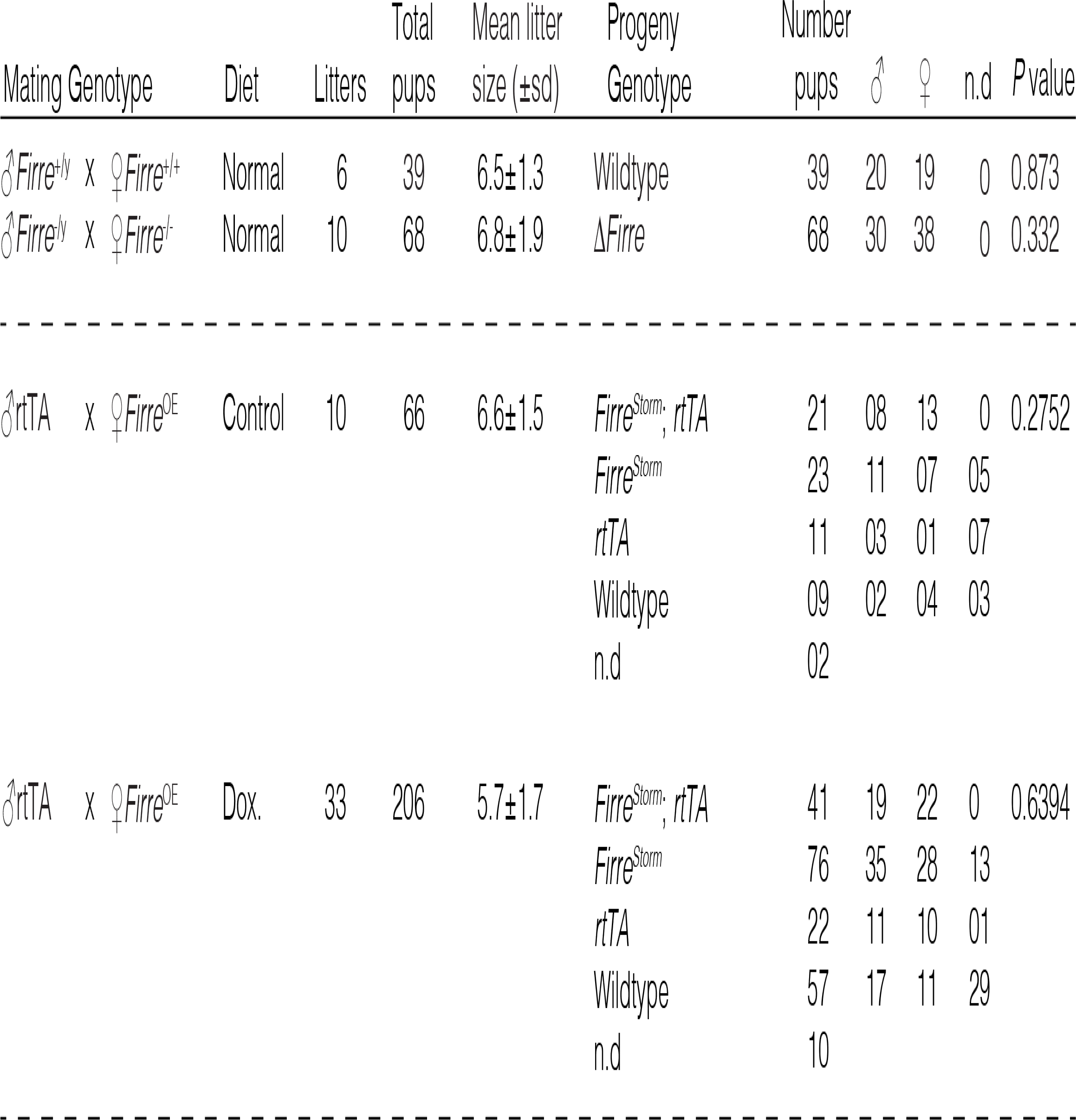
Genotype and male and female distribution in Δ*Firre* and *Firre* overexpressing mice. Genotyping from progeny at P7 from intercrosses between male WT and female WT; male Δ*Firre* and female Δ*Firre*; male rtTA and female *Firre*^OE^ no dox; and male rtTA and female *Firre*^OE^ dox-diet mice. Litter size shown as mean with standard deviation (s.d.), not determined (n.d.), Chi-square statistic reported (p-value).

**Extended Data Table 2. Differential gene expression in midbrain tissue from E11.5 wild-type and Δ*Firre* embryos.**

**Extended Data Table 3. Differential gene expression in forebrain tissue from E11.5 wild-type and Δ*Firre* embryos.**

**Extended Data Table 4. Differential gene expression in presomitic mesoderm tissue from E11.5 wild-type and Δ*Firre* embryos.**

**Extended Data Table 5. Differential gene expression in lung tissue from E11.5 wild-type and Δ*Firre* embryos.**

**Extended Data Table 6. Differential gene expression in hindlimb tissue from E11.5 wild-type and Δ*Firre* embryos.**

**Extended Data Table 7. Differential gene expression in forelimb tissue from E11.5 wild-type and Δ*Firre* embryos.**

**Extended Data Table 8. Differential gene expression in liver tissue from E11.5 wild-type and Δ*Firre* embryos.**

**Extended Data Table 9. Differential gene expression in heart tissue from E11.5 wild-type and Δ*Firre* embryos.**

**Extended Data Table 10. Differential gene expression in male CLPs from wild-type and Δ*Firre*.**

**Extended Data Table 11. Differential gene expression in male CLPs from ΔFirre and Δ*Firre; Firre^rescue^*.**

